# The regulatory landscape of early maize inflorescence development

**DOI:** 10.1101/870378

**Authors:** Rajiv K. Parvathaneni, Edoardo Bertolini, Md Shamimuzzaman, Daniel Vera, Pei-Yau Lung, Brian R. Rice, Jinfeng Zhang, Patrick J. Brown, Alexander E. Lipka, Hank W. Bass, Andrea L. Eveland

**Author notes:** These authors contributed equally to this work (Rajiv K. Parvathaneni, Edoardo Bertolini). USDA-ARS Edward T. Schafer Agricultural Research Center, Fargo, ND 58102. Department of Genetics, Harvard Medical School, Boston, MA 02115.

## Abstract

**Background:** The functional genome of agronomically important plant species remains largely unexplored, yet presents a virtually untapped resource for targeted crop improvement. Functional elements of regulatory DNA revealed through profiles of chromatin accessibility can be harnessed for fine-tuning gene expression to optimal phenotypes in specific environments.

**Result:** Here, we investigate the non-coding regulatory space in the maize *(Zea mays)* genome during early reproductive development of pollen- and grain-bearing inflorescences. Using an assay for differential sensitivity of chromatin to micrococcal nuclease (MNase) digestion, we profile accessible chromatin and nucleosome occupancy in these largely undifferentiated tissues and classify at least 1.6 percent of the genome as accessible, with the majority of MNase hypersensitive sites marking proximal promoters, but also 3’ ends of maize genes. This approach maps regulatory elements to footprint-level resolution. Integration of complementary transcriptome profiles and transcription factor occupancy data are used to annotate regulatory factors, such as combinatorial transcription factor binding motifs and long non-coding RNAs, that potentially contribute to organogenesis, including tissue-specific regulation between male and female inflorescence structures. Finally, genome-wide association studies for inflorescence architecture traits based solely on functional regions delineated by MNase hypersensitivity reveals new SNP-trait associations in known regulators of inflorescence development as well as new candidates.

**Conclusions:** These analyses provide a comprehensive look into the cis-regulatory landscape during inflorescence differentiation in a major cereal crop, which ultimately shapes architecture and influences yield potential.

## Background

Growth and development of all organisms depends on coordinated regulation of gene expression in time and space. In order for this to occur, chromatin conformation must be remodeled accordingly so that specific sequences in the DNA are accessible to regulatory transcription factors (TFs) [1]. Eukaryotic chromatin is packaged into arrays of nucleosomes, which each consist of 147 bp of DNA wrapped around a histone octamer. Dynamic shifts in nucleosome positioning affect gene regulation by modifying availability of cis-regulatory elements (CREs) to which combinations of TFs bind in a spatiotemporal manner [2–4]. Genetic variation in CREs and tissue specificity of regulatory factors can contribute to phenotypic plasticity. Since CREs are difficult to identify by sequence alone, various Encyclopedia of DNA Elements (ENCODE)-like projects have emerged with the goal of annotating the non-coding, functional genome in a given species by generating spatiotemporal maps of chromatin accessibility, TF occupancy, protein and DNA modifications, and gene expression [5,6]. Although similar datasets are beginning to mature for a number of plant species [7–10], our knowledge of gene regulation in plants remains limited, and notably in our most economically important crops [11].

Various methods have been used to map genome-wide chromatin accessibility and nucleosome occupancy [12–15] and their use is based on the idea that TF occupancy occurs in nucleosome-free regions that present biochemically as nuclease accessible or hypersensitive (HS). Enzymes with nuclease activity, such as deoxyribonuclease I (DNaseI) and micrococcal nuclease (MNase), have thus been used to map nuclease HS regions of the chromatin as a proxy for regulatory sequences in plants [16–20]. Overlaying HS regions with TF binding and/or haplotype maps have identified genetic loci that contribute, for example, to human disease [21,22]. Since the vast majority of genetic variants map to noncoding intergenic and intronic regions, chromatin accessibility provides a filter to enrich for potential functional variants and facilitate discovery of new regulatory loci. In maize, it was reported that approximately 1% of the genome was accessible based on differential MNase sensitivity in nuclei from young seedlings, yet this fraction explained 40% of the heritable variation [17]. These results indicated that, as in animal systems, functional polymorphisms are enriched in accessible regions of plant genomes, which has significant implications for advancing genome-assisted breeding.

Based on studies in animals and plants, CREs can reside within the core or proximal promoters of genes (i.e., at or near the transcriptional start site (TSS)), or within cis-regulatory modules distal to their target genes, such as enhancer sequences that can influence gene expression at long range [2,3]. Functional studies on distal enhancers, predominantly in animal systems, showed they can integrate various signals and fine-tune gene expression through combinatorial recruitment of TFs that are available in distinct spatiotemporal patterns [23–25]. Aside from a handful of functionally tested cases, the majority identified by Quantitative Trait Loci (QTL), our knowledge of distal regulatory sequences in plants remains poorly defined [26]. In maize, the domestication genes *teosinte branched1 (tb1)* and *grassy tillers 1 (gt1)* are regulated by distal enhancer sequences. Genetic variation in an enhancer region ~60 kb upstream of *tb1* modulates its expression through two distinct transposable elements (TEs) [27,28]. Similarly, variation in a region ~7.5 kb upstream of *gt1* controls its expression [29]. Genome-wide surveys of putative enhancer regions have been carried out in Arabidopsis, rice and maize [7,16,30,31], yet very few have been functionally validated.

Maize is the cereal crop with the largest dollar value in the U.S. and abroad. Non-coding sequence makes up ~98% of the maize genome, the vast majority of which remains unexplored. In this study, we used a method that leverages differential sensitivity of chromatin to various concentrations of MNase followed by high-throughput sequencing (MNase-seq) to map global chromatin accessibility and nucleosome occupancy in maize inflorescence primordia early in development. Maize forms male and female flowers on separate inflorescences; the male tassel is formed apically following the transition from vegetative development and female ears are produced in the axils of leaves. Tassel and ear share common developmental programs and are morphologically similar at the early primordia stages. But variations on these developmental programs at the molecular level result in distinct structural features at maturity that directly affect yield potential, e.g., outgrowth of branches in the tassel and development of kernel rows on the ear [32,33].

Our analysis of tassel and ear specifically examines primordia, which are enriched for meristematic cells ideal for mapping of HS signatures to footprint-level resolution. These accessible chromatin maps were integrated with complementary genomics datasets from comparable tissues, e.g., transcriptome profiles and TF occupancy maps, to validate functional regions in the maize genome. We also annotated a suite of long non-coding (lnc)RNAs associated with signatures of chromatin accessibility in the young maize inflorescences, which showed spatiotemporal specificity. Finally, we identified novel trait-SNP associations for key inflorescence architecture traits by using the functional genome to guide GWAS analyses. Results from this work provide a foundation for deeper explorations of the regulatory mechanisms underlying early developmental transitions during maize tassel and ear morphogenesis, which can be leveraged for crop improvement through molecular breeding and/or precision engineering.

## Results

### Mapping accessible chromatin in early development of maize inflorescences

To determine the regulatory landscape of the maize genome during early inflorescence development, we generated genome-wide, tissue-specific maps of accessible chromatin from immature tassels and ears. We used a method that defines differential sensitivity of chromatin using different concentrations of micrococcal nuclease (MNase), which was previously demonstrated in maize for vegetative tissue types [17,19,34]. Here, maize tassel and ear primordia were precisely staged and hand-dissected from greenhouse-grown, B73 inbred maize plants (Additional file 1: Figure S1). For each tissue, approximately 170 individual primordia (1-5 mm in size) were pooled for each of two biological replicates. This developmental sampling scheme is comparable to previous genomics analyses for these tissues [32,35,36]. From each sample, fixed, intact nuclei were isolated and divided into two pools for “light” and “heavy” MNase digestion, and genomic library construction (Additional file 1: Figure S1).

Mapped reads from light and heavy digests were compared to reveal regions of differential nuclease sensitivity (DNS), including MNase sensitive (accessible chromatin) and MNase resistant (closed chromatin) signatures (Additional file 1: Table S1). Since the exonuclease activity of MNase results in protected fragments of various sizes, data were analyzed in different ways relative to the DNA fragment sizes. For example, nucleosome-protected fragments are expected to be greater than 130 bp in length, whereas smaller fragments (< 131 bp), particularly those resulting from a light MNase digest, are likely associated with small, sub-nucleosomal particles in chromatin, biochemically defined as open and accessible. Therefore, selective mapping of different size classes from the light and/or heavy digests can provide different views of the chromatin landscape, as previously described [15,37]. We mapped read coverage across the maize genome for: i) all size classes from light and heavy digests (provides information on accessible chromatin as well as nucleosome occupancy); ii) DNS analysis; and iii) small fragment (< 131 bp) coverage from the light digests only, hereafter referred to as Light digest Coverage Small fragments (LCS) (Figure 1a). Overall, MNase hypersensitivity (HS) associated with gene-rich regions of the genome and MNase resistance was largely associated with repetitive sequence.

**Figure 1.**
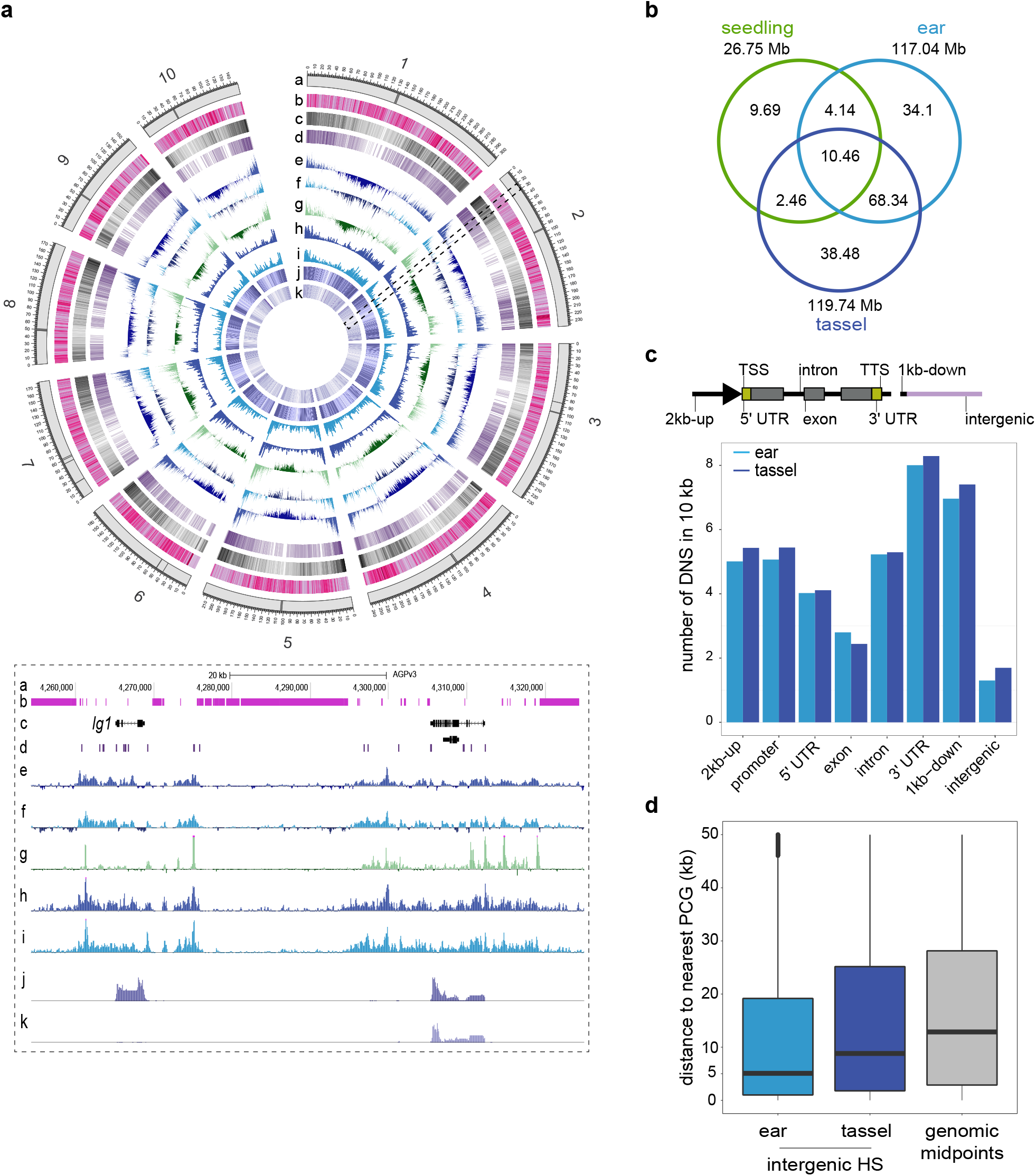
Genome-wide mapping of chromatin accessibility in early tassel and ear primordia. **(a)** Genome-wide visualization of MNase HS in tassel and ear and aligned against other genomic features: a – Chromosome ideograms; black bars indicate centromeres, b – Repeat elements, c – Gene density, d – Density of conserved non-coding sequences (CNSs), e – Differential nuclease sensitivity (DNS) from tassel, f – DNS from ear, g – DNS from seedling shoot, h – Density of small fragments (<131bp) from the light MNase digest (LCS) in tassel, i – LCS in ear, j – Density of gene expression from RNA-seq in 1-2 mm tassel primordia, k – Density of gene expression from RNA-seq in 1-2 mm ear primordia. A 60 kb slice on chromosome 2 is represented in a genome browser view below and includes the *liguleless 1* gene and a prominent HS region approximately 20 kb upstream. **(b)** The venn diagram compares the portion of MNase HS genome (in Megabases) based on DNS in tassel, ear and shoot tissues. **(c)** Distribution of MNase HS sites (based on peak midpoint) across eight genomic features relative to protein coding genes: within 1 and 2 kb upstream of the TSS, the proximal promoter defined as 1 kb upstream and 200 bp downstream of the TSS, 5’ UTR, exon, intron, 3’ UTR, within 1 kb downstream of the TTS, intergenic. Density of DNS signatures were normalized per 10 kb of genomic space per feature. In both tassel and ear, the promoter and 3’ regions of genes are most accessible. **(d)** Distribution of distances (in the range of 50 kb) from the midpoints of intergenic MNase HS sites to the closest protein coding gene in ear and tassel. The grey bar plots the distribution of midpoints of genomic space between any two sequential genes in the genome.

To classify genomic regions of chromatin accessibility, we used the peak-calling algorithm iSeg [38] (see Methods; Additional file 1: Figures S2 and S3, Tables S2 and S3; Additional files 2-5: Datasets S1-S4). Based on genomic overlap of HS regions, approximately 67.3% of the accessible genome was shared between tassel and ear (Figure 1b). If considering genomic space within 2 kb of genes, this increases to 75%. The high concordance between tissues was expected given morphological similarities between tassel and ear at the early stage examined here. Approximately 33% of tassel accessible regions were found only in tassel, and 26% were specific to ear (Additional file 1: Figure S4; Additional file 6: Dataset S5). We also re-analyzed publicly available MNase-seq data from arial maize seedling tissues [17] using iSeg (Figure 1a-b). As was reported previously, 1.07% of the genome was MNase HS in the seedling shoot. Approximately 51% of MNase HS in seedling shoot was shared in inflorescence primordia (Figure 1b).

We next determined the genomic distribution of MNase HS by mapping the midpoint of each region to genic and intergenic features (Figure 1c; Additional file 1: Table S4). We defined the proximal promoter of protein coding genes (PCGs) to include 1 kb upstream and 200 bp downstream from the TSS, which showed higher peak densities compared to within gene body features. Strikingly, high densities of HS were also observed in the 3’ UTR and downstream regions flanking genes (Figure 1c). While DNS profiles showed similar peak intensities at the TSS and Transcriptional Termination Site (TTS) of genes, densities of LCS were higher at TSSs (Additional file 1: Figure S5). This indicates that the promoter region of genes is characterized by more non-nucleosomal footprints while downstream flanks are predominantly nucleosome-bound. For HS signatures that mapped to intergenic regions, the distance to the nearest PCG was determined. Intergenic HS regions were positioned significantly closer to genes than by random chance in both tassel and ear, however the median distance to genes was significantly smaller in ear (~5 kb) than tassel (~10 kb; Welch’s *t-test*(p-value < 2.2 e^-16^)) (Figure 1d). Mapping tassel- and ear-specific HS sites accentuated the differences observed in their genomic distributions, with tassel HS enriched in intergenic regions, and ear-specific HS more prominent in gene body features (Additional file 1: Figure S4, Table S4).

We asked whether the increased tassel-specific accessibility in the intergenic space was associated with transposable elements (TEs). Approximately 85% of the maize genome is made up of TEs, and 50% of the transposon component resides within gene-rich regions of the genome [39,40].

Based on TE annotations in the maize genome [41], we found enrichment of MNase HS associated with TEs relative to randomized intergenic regions (*p* value < 0.001, permutations *n* = 1,000), as previously shown for other tissue types [16,42]. Notably, there was increased chromatin accessibility associated with retrotransposons in tassel primordia compared to ear (*p* value < 0.001, permutations *n* = 1,000; Additional file 1: Figure S6). This is consistent with early activation of TEs in the male germline as it was recently shown that in tassel primordia, TEs are activated much earlier than was described for Arabidopsis, well before the development of male-specific reproductive organs [43].

### Differential HS in gene promoters reveals regulatory DNA associated with organ-specific expression

Previous work comparing chromatin accessibility in different maize tissues showed that HS in proximal promoters of genes tended to correlate with gene expression [17,19]. Here, we leveraged publicly available RNA-seq datasets from early tassel and ear primordia comparable with those used for our MNase-seq data [32]. PCGs were divided into three equal groups based on their normalized expression levels in Transcript Per kilobase Million (TPM), and for each group (none-low, mid, and high expression; Additional file 7: Dataset S6), MNase HS was plotted with respect to a consensus promoter region (± 1 kb around the TSS; Figure 2a). Enrichment of MNase HS was observed in proximal promoters of expressed genes in both tassel and ear, indicating a positive correlation between degree of chromatin accessibility and gene expression level (Figure 2a; Additional file 1: Figure S7).

**Figure 2.**
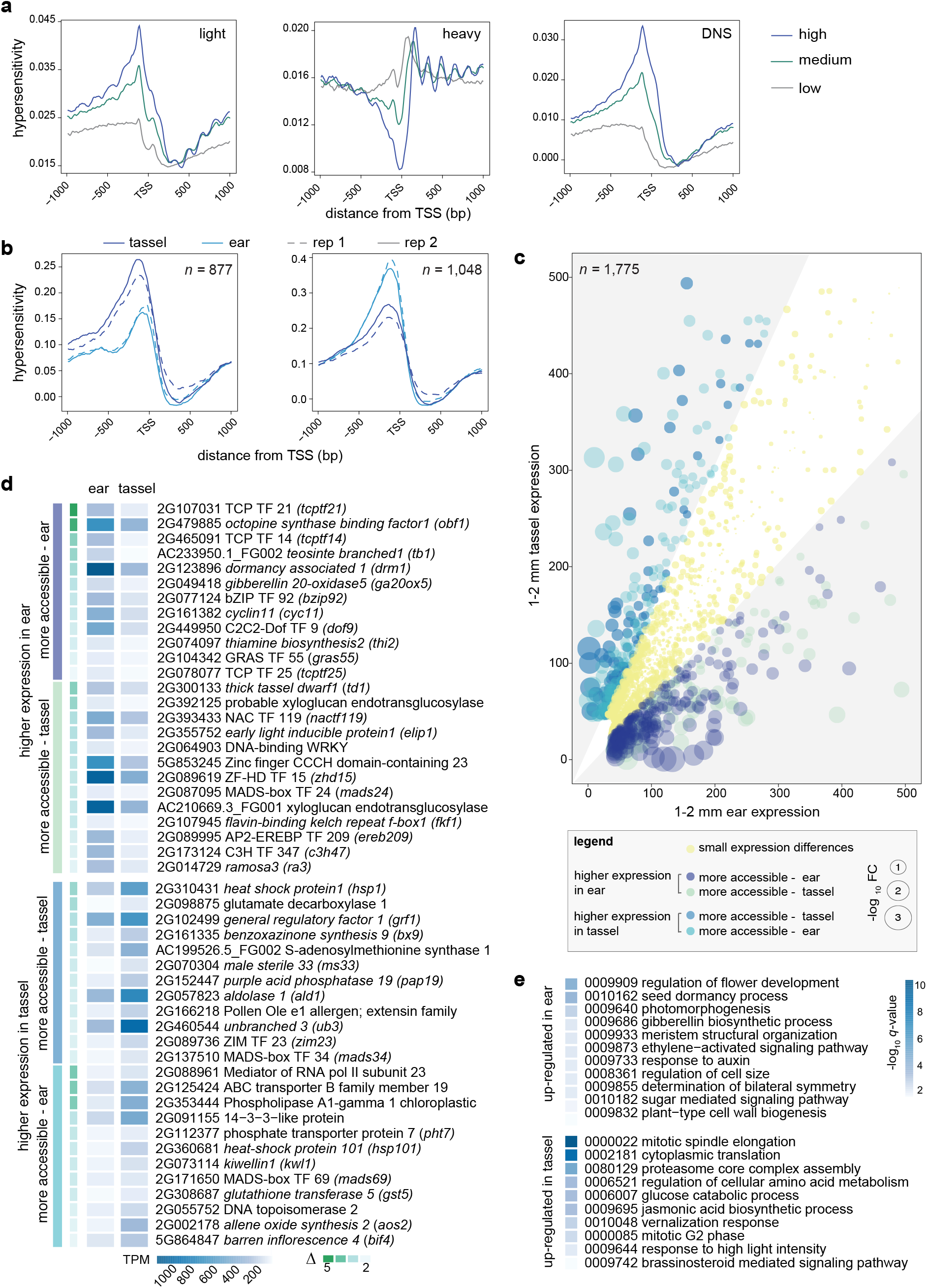
Differential MNase HS and expression changes between tassel and ear. **(a)** MNase HS relative to the TSS of three expression classes of PCGs in tassel. HS is plotted as average read density from the light, heavy and differential (light-heavy) MNase analysis ± 1,000 bp surrounding the TSS of a composite PCG model divided into 3 tertiles (high, medium and low) based on their TPM values. Results from ear are similar and presented in Figure S7. **(b)** Distribution of MNase HS around the TSS of 1,925 PCGs that were differentially accessible within 500 bp upstream of the TSS (877 showed increased HS in tassel and 1,048 in ear). **(c)** Expression profiles of differentially accessible PCGs in tassel compared to ear primordia. Genes are represented as colored bubbles of different sizes (log_10_ fold change of gene expression between the two tissues). Bubble color indicates one of five gene classes as indicated in the legend. Grey areas in the scatterplot represent genes with a fold change difference greater than |1.5|. Genes with expression levels of > 500 TPM (n=150) were excluded from the visualization. **(d)** A subset of differentially regulated PCGs and their expression profiles are grouped in four classes: more accessible and higher expression in tassel, more accessible and higher expression in ear, more accessible and lower expression in tassel, and more accessible and lower expression in ear. Genes are listed in descending order by their delta value, which represents the degree of differential accessibility in the immediate proximal promoter. Functional annotations are based on information at MaizeGBD. (e) Subset of GO categories overrepresented among differentially accessible PCGs that showed gene expression changes between tassel and ear.

We expect chromatin accessibility to be highly dynamic during meristem development and organogenesis. Genes that control meristem determinacy are largely expressed in both tassel and ear, but it is the precise regulation of these genes that fine-tune developmental processes that lead to subtle morphological differences. By defining differential promoter accessibility and/or usage coupled to gene expression changes, we hope to resolve tissue-specific regulatory differences that underlie variation in branching, or meristem determinacy, between tassel and ear. We asked whether differential degrees of HS in proximal promoters of genes between tassel and ear translated to differential expression. Based on a set of criteria described in the Methods, a differential DNS analysis was performed for promoters of PCGs (500 bp directly upstream of the TSS). This revealed 1,925 genes with differential accessibility in their immediate proximal promoters; 877 showed increased HS in tassel and 1,048 in ear (Figure 2b; Additional file 8: Dataset S7). Expression levels of these genes were plotted (tassel relative to ear), and classified based on the degree and direction of expression change with respect to promoter accessibility (Figure 2c; Additional file 8: Dataset S7). Approximately 41% (794) showed substantial expression differences between tassel and ear, including direct correlations with chromatin accessibility as well as anti-correlated trends (Figure 2c). The latter could be indicative of repressor binding and this was observed for approximately half of the genes. The rest showed little differences in expression between organs despite the differential accessibility in the immediate promoter and could reflect poised transcriptional complexes, for example.

Among the differentially regulated genes (i.e., differential accessibility in the proximal promoter and altered gene expression between tassel and ear) were 112 TFs, including multiple members of TF families implicated in meristem development and organogenesis (Figure 2d; Additional file 1: Figure S8). The majority of these differentially regulated TFs were more highly expressed in ear primordia, which were enriched for members of ZF-HD (q = 7.67e-04), GRF (*q* = 1.19e-02), GRAS *(q* = 4.19e-02), ERF *(q* = 5.67e-02) and TCP (Teosinte branched/Cycloidea/PCF; *q* = 5.67e-02) families. The latter included five differentially regulated family members including *tb1* (AC233950.1_FG002), a class II TCP that is well known for its function in repressing shoot branching in maize [44]. Consistent with a function for *tb1* in repressing branching in ears, there was increased accessibility in its promoter and much higher expression in ear primordia (Figure 2d). The other four TCPs (GRMZM2G078077, GRMZM2G107031, GRMZM2G445944, GRMZM2G465091) are of the class I type of TCP TFs. Class I TCPs have been generally less studied for their roles in development, however, several members have been shown to control aspects of cell division and differentiation [45]. All five TCPs showed strikingly higher expression in ears and notable increases in promoter accessibility.

We performed functional enrichment analysis on differentially regulated genes using Gene Ontologies (GO) based on maize-Gamer [46] annotations. Functional categories overrepresented among genes that were up-regulated in ear primordia included biological processes related to flower development and organogenesis (e.g., GO:0009909: regulation of flower development; *q* = 4.1e-05, GO:0009933: meristem structural organization; *q* = 8.73e-03, GO:0008361: regulation of cell size; *q* = 1.12e-02) (Figure 2e; Additional file 8: Dataset S7). In addition to tb1, other classical maize genes known to repress branching and/or promote meristem determinacy were in this class, e.g. *dormancy associated1 (drm1), ramosa 3 (ra3)* [47] and *thick tassel dwarf (td1)* [48] (Figure 2d). Increased transcription of these genes is consistent with a determinacy program being imposed on axillary meristems in the ear compared to tassel; the latter produces indeterminate branch meristems at this early stage of development.

In contrast, differentially regulated genes with higher expression in tassel primordia tended to be involved in cell growth processes including translation (GO:0002181: cytoplasmic translation; *q* = 6.26e-10), protein synthesis and turnover, and sugar metabolism (GO:0006007: glucose catabolic process; *q* = 4.4e-04) (Figure 2d,e; Additional file 8: Dataset S7). This could be related to increased protein and energy metabolism required for outgrowth of indeterminate branch meristems. Interestingly, distinct hormone pathways appear to be differentially regulated in tassel and ear. Auxin, ethylene and GA-related pathway components tend to be up-regulated in ear, while genes involved in JA and BR synthesis and signaling are up-regulated in tassel (Figure 2d,e).

### MNase HS signatures predict TF binding sites to footprint level resolution

To test whether HS sites called from LCS are a proxy for TF binding, we overlaid ChIP-seq data generated from comparable inflorescence tissue for two developmental regulators; KNOTTED 1 (KN1), a homeodomain TF that regulates maintenance of all plant meristems [36], and FASCIATED EAR 4 (FEA4), a bZIP TF that regulates meristem size [35]. More than 90% of KN1 and FEA4 high confidence ChIP-seq peaks overlapped with the MNase HS sites, and approximately 97% of sites co-bound by KN1 and FEA4 were accessible (Figure 3a; Additional file 1: Figure S9). It was previously shown that KN1 and FEA4 TFs co-occupy many sites during early maize inflorescence development to regulate lateral organ initiation [35] and we expect co-bound regions to be regulatory. Genes co-bound by KN1 and FEA4 in their proximal promoters showed stronger and broader MNase HS signatures than those bound by either TF alone, consistent with highly accessible promoters occupied by multiple TFs (Figure 3b and Additional file 1: Figure S9; Additional File 9: Dataset S8). Accessible intergenic sites that are co-bound by these TFs are potential long-range transcriptional regulators. For example, FEA4 and KN1 co-bound an accessible region that aligned with a known enhancer element ~65 kb upstream of *tb1* (Figure 3c). Another co-bound region 6 kb upstream of the *gnarley 1 (gn1)* homeobox locus was accessible only in inflorescence tissue (Figure 3c).

**Figure 3.**
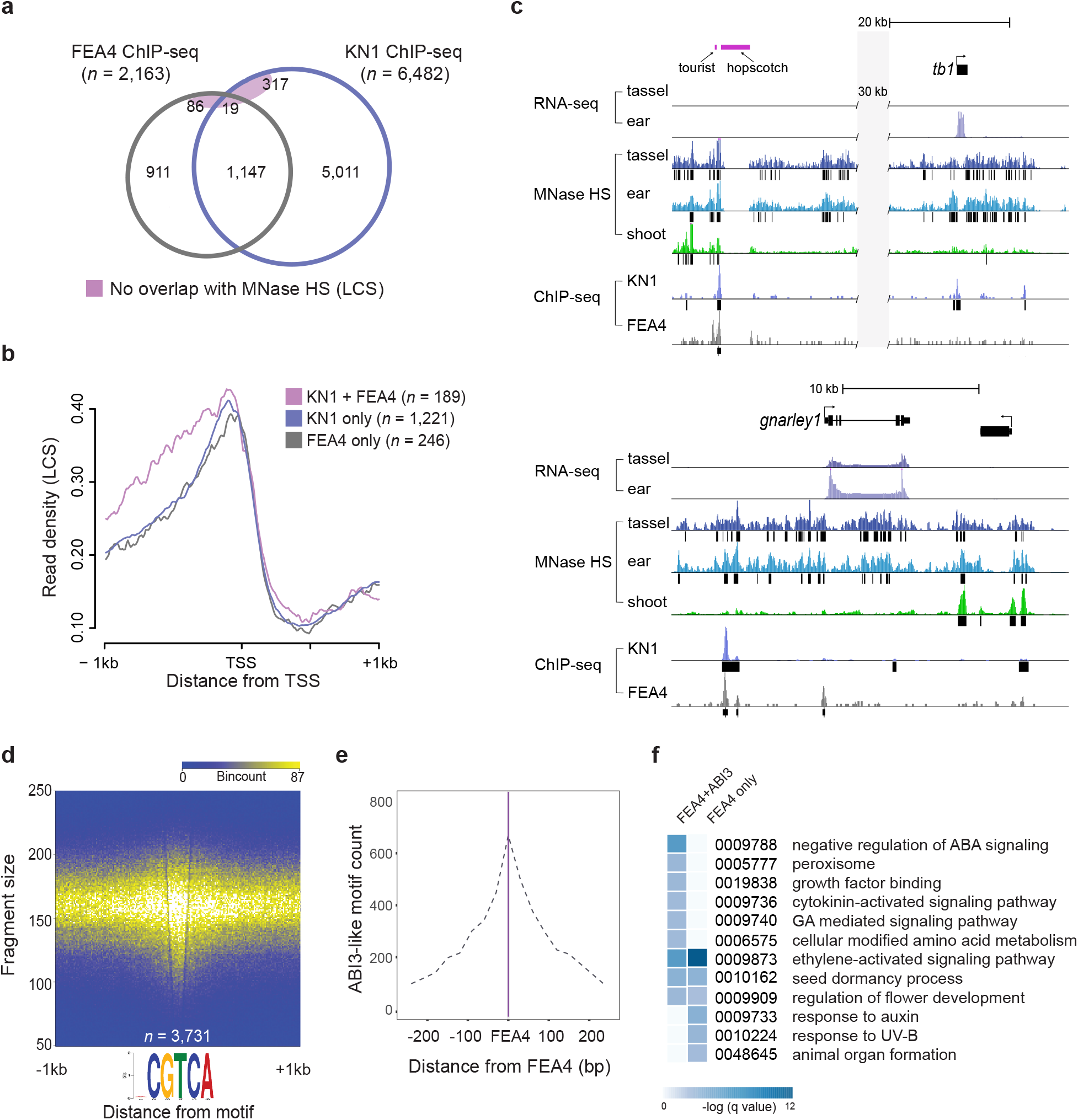
MNase HS as a proxy for TF-DNA binding. **(a)** Overlap of high confidence peaks from FEA4 and KN1 ChIP-seq experiments with MNase HS regions based on LCS data. Less than two percent of co-bound sites and six percent of all sites do not overlap MNase HS. **(b)** Read density (reads per million) of LCS are plotted around a consensus TSS of genes bound by KN1 and/or FEA4. Genes cobound by both TFs in the proximal promoter show a wider range of accessibility. Data are shown for tassel; similar profiles are shown for ear presented in Figure S9. **(c)** Examples of intergenic MNase HS sites in tassel, ear, and/or shoot that are co-bound by KN 1 and FEA4: a known enhancer element 65 kb upstream of *tb1,* and a site 6 kb upstream of the *gnarley 1 (gn1)* locus. (d) Fragment densities from the light MNase digest plotted around high-confidence FEA4 ChIP-seq peaks show strong enrichment at the consensus FEA4 binding motif, **(e)** Density of ABI3-like motifs were plotted in relation to experimentally validated FEA4 binding sites. The ABI3-like motif was discovered by mining HS regions called by iSeg that overlapped consensus FEA4 binding sites. ABI3-like motifs are most densely positioned within 50 bp of FEA4 motifs. **(f)** GO terms overrepresented among genes within 1 kb of FEA4-ABI3-like motif modules (ABI3-like motifs within 50 bp of FEA binding sites) and compared with genes within 1 kb of random FEA4 motifs of equal number.

To further test whether density of MNase small fragments could locate TF binding sites, we mined sequences under 2,163 high-confidence FEA4 ChIP-seq peaks that overlapped with MNase HS. We resolved a consensus FEA4 motif (NCGTCA; *p* = 1.3e-183). Density plots of LCS showed high enrichment centered around FEA4 motifs in both tassel and ear (Figure 3d; Additional file 1: Figure S9). We next used the iSeg-defined regions overlapping the FEA4 motifs to computationally screen for other known TF binding motifs enriched around FEA4 binding (Additional file 1: Figure S10). Multiple Em for Motif Elicitation (MEME) was used to identify *de novo* motifs, which were compared to experimentally validated plant Position Weight Matrices (PWMs) to define best matches. In addition to the FEA4 (bZIP) motif itself and KN1, we identified a highly overrepresented motif that matched an experimentally validated binding site for a plant B3-domain TF, Abscisic Acid Insensitive (ABI) 3, based on two different motif scanning methods (Additional file 1: Figure S10). This motif was present within 70% of the FEA4 binding site regions. Distribution of this motif relative to FEA4 binding showed strong enrichment within 50 bp (Figure 3e; Additional file 10: Dataset S9). Functional enrichment analysis of genes proximal (within 1kb) to FEA4-ABI3-like cis-element modules (the two motifs within 50 bp of each other) showed overrepresentation of GO terms related to ABA-related signaling, e.g., negative regulation of abscisic acid-activated signaling pathway (GO:0009788; *q* = 1.84e-03) and seed dormancy process (GO:0010162; *q* = 0.02), and other hormone pathways such as ethylene activated signaling (GO:0009873; *q* = 0.002) compared to genes flanking a random set (n = 1039) of FEA4 motifs (Figure 3f; Additional file 10: Dataset S9). This suggests potential functional significance of a FEA4 regulatory interaction with an ABI3-like B3-domain TF to control a particular suite of genes, possibly in a combinatorial way.

The average size of HS regions called from the small fragment data was 165 bp. We asked whether we could refine putative TF footprints within these regions by mapping just the midpoints of the LCS. Theoretically, the mid-point of these read fragments would be most protected from MNase activity by a TF binding to DNA. We used iSeg to call footprints based on read density at midpoints. At a stringent bc 4.0 threshold, an average of ~350,000 footprints were called in tassel and ear with a median size of 21 bp (Additional file 11: Dataset S10), which constituted less than 0.01% of the maize genome. Alignment with FEA4 binding sites was statistically significant (*p* = 0.001, 1,000 iterations; Additional file 1: Figure S11).

We next performed motif enrichment analysis within the footprints to predict candidate TF binding that drives suites of differentially regulated genes between tassel and ear primordia (Figure 2c). For genes that showed an expression change between tissues and increased promoter accessibility in tassel or ear, footprints within 2 kb upstream and 200 bp downstream of the TSS, and 1 kb downstream of the TTS, were mined for *de novo* motifs using Discriminative Regular Expression Motif Elicitation (DREME) [49]. PWMs of highly enriched motifs were matched against the JASPAR database of known plant TF binding sites (http://jaspar.genereg.net/). Genes that showed increased promoter accessibility in tassel were highly enriched for ERF and ARF TF motifs, while those that were more accessible in ear were enriched for WRKY, C2H2 zinc finger, TCP and MYC elements (Figure 4; Additional file 1: Table S5). The five TCP TFs that were up-regulated in ear primordia (Figure 2d) all had multiple occurrences of the predicted TCP motif (RGCCS), suggesting regulation of individual TCPs by each other and/or other family members (Figure 4a-b). We also scanned all mapped footprints in tassel and ear for the 21 experimentally validated TCP PWMs that exist in JASPAR plant (Additional file 1: Table S6). We found TCP binding sites overall were more enriched in ear HS footprints compared to tassel, and that PWMs associated with TCP23 and TCP15 from Arabidopsis were most highly enriched.

**Figure 4.**
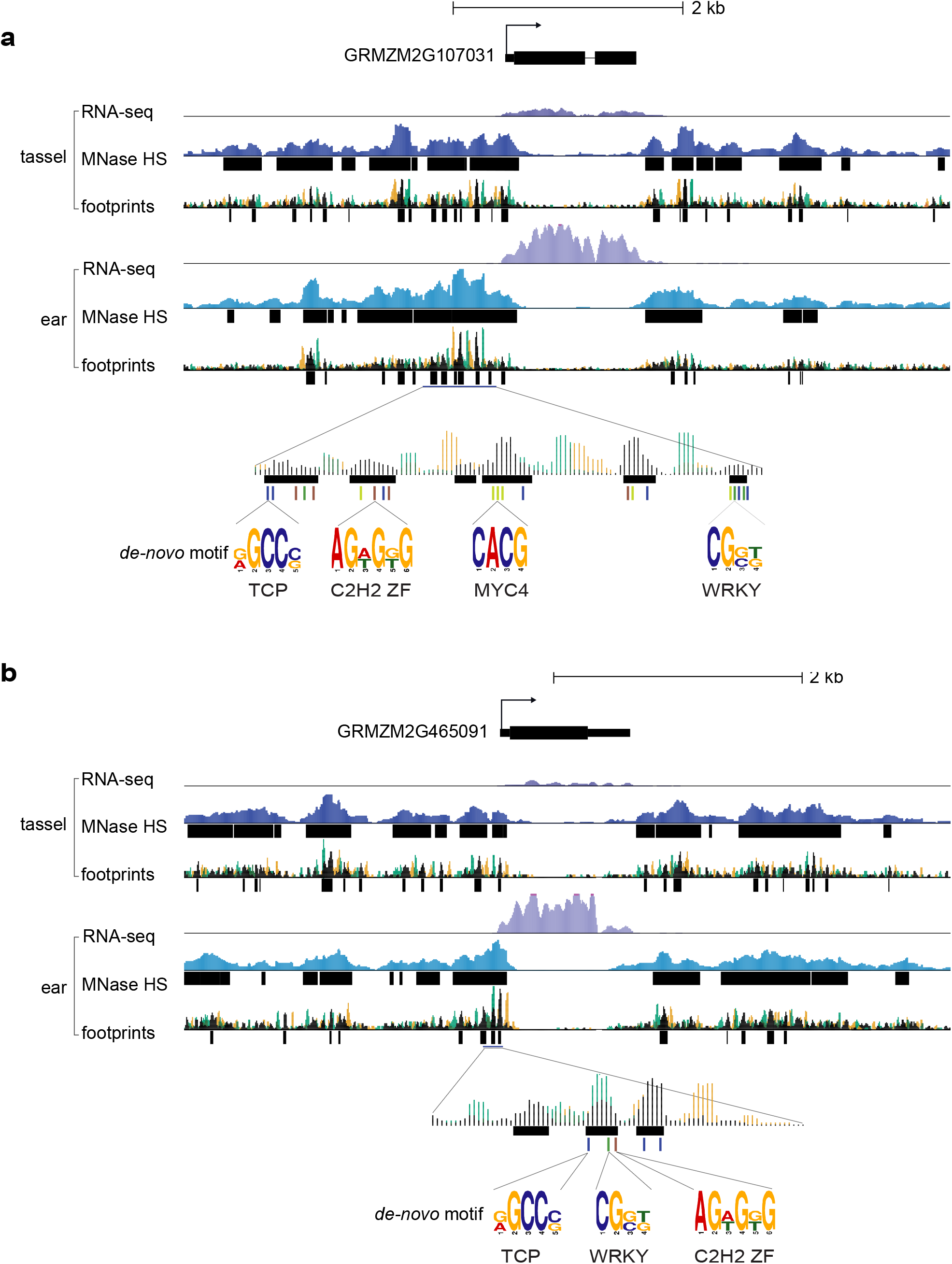
TCP TFs up-regulated in ear primordia harbor TCP binding motifs in promoter footprints. Genomic regions are shown surrounding two class I TCP TFs that were up-regulated in ear primordia compared to tassel. For (a) GRMZM2G107031 and (b) GRMZM2G465091, RNA-seq data show enhanced expression in ear, with increased MNase HS. MNase footprint regions were called by mapping fragment center hotspots, shown in black. The 5’ (green) and 3’ (orange) fragments ends are also indicated. Four motifs (TCP, C2H2. MYC4, WRKY) were significantly enriched in footprints in promoters of genes up-regulated in ear primordia and are color-coded.

### Long non-coding RNAs associate with accessible chromatin in inflorescence primordia and mark putative distal regulators

In addition to enrichment of MNase HS at the proximal 5’ and 3’ ends of PCGs, we observed numerous intergenic accessible regions greater than 2 kb away, but largely within 10 kb of genes (Figure 1d). Based on our knowledge of gene regulation in other systems, at least some of these distal accessible regions may mark enhancers or other long-range regulators of gene expression. Reports in both animal and plant systems have linked enhancers with transcription of long non-coding RNAs (lncRNAs), which have been shown to control distal gene expression in various ways [50–52]; e.g., aiding chromatin looping to modulate expression of adjacent or distant PCGs [53–55].

Since lncRNAs are highly context-specific, we cataloged the spatiotemporal expression of lncRNAs in maize inflorescence primordia to i) investigate their association with intergenic HS chromatin signatures and ii) to extend our regulatory maps for tassel and ear development. We used a pipeline described by De Quattro et al. (2017) [56] to annotate these lncRNAs by re-analyzing RNA-seq data from a developmental series of tassel and ear primordia stages [32]. We identified 2,679 high-confidence lncRNAs derived from 2,520 unique loci (Figure 5a; Additional file 1: Figures S12 and S13; Additional file 12: Dataset S11; Additional file 13: Dataset S12). These lncRNAs showed spatiotemporal expression differences during tassel and ear development, with expression generally most prominent in undifferentiated inflorescence meristem (IM) tissue at the earliest stages of development (Figure 5a). We used the Shannon Entropy (SH) calculation [57] to determine spatiotemporal specificity of lncRNA expression across early tassel and ear development (Additional file 1: Figure S13). A subset of lncRNAs (*n* = 52) was specifically expressed in a particular meristem type or developmental stage (SH ≥ 0.6; Additional file 1: Figure S14), while the majority were dynamically expressed throughout inflorescence development (SH ≤ 0.2).

**Figure 5.**
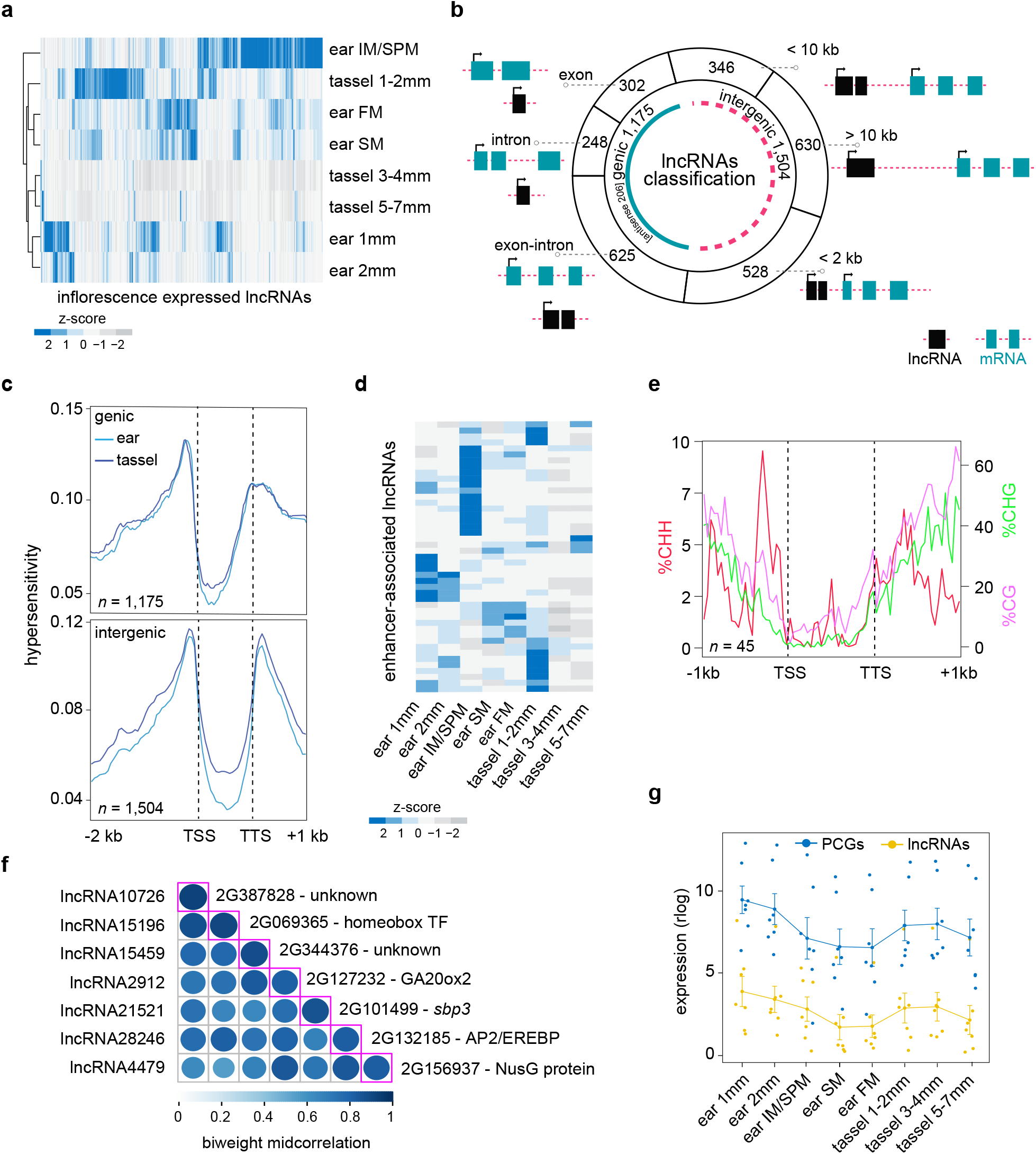
A catalog of long non-coding (lnc)RNAs expression during early maize inflorescence development. **(a)** Expression profiles (plotted as normalized z-score) for 2,679 lncRNAs across different stages of ear and tassel development. Euclidean distance was computed across rows (lncRNAs) and columns (tassel and ear developmental stages), and data were hierarchically clustered using the maximum linkage method. **(b)** The various classifications of lncRNAs used in this study, including their positions relative to PCGs. **(c)** MNase HS profiles across a composite genic and intergenic lncRNA model show highest densities around the TSS and TTS, similar to PCGs. **(d)** Expression profiles (plotted as normalized z-scores) during early ear and tassel development of lncRNAs that overlapped putative enhancer loci identified from maize leaf and husk tissue (enhancer-associated intergenic lncRNAs) [16]. **(e)** Context-specific methylation profiles plotted across a composite lncRNA model based on 45 enhancer-associated intergenic lncRNAs. CHH (red) methylation is plotted on the left, and CG (purple) and CHG (green) methylation on the right. **(e)** A representative co-expression module of lncRNA and PCG pairs located in a genomic window of < 2 kb. The diagonal axis indicates the co-expression value between each lncRNA and PCG pair expressed as the biweight midcorrelation. **(f)** Expression values of the seven lncRNA-PCG pairs within the module in **(e)**. Dots represent the rlog values of lncRNAs (yellow) and PCGs (blue), and the line indicates the average expression level.

LncRNAs were classified as genic or intergenic. Based on overlap with known gene features, genic lncRNAs were further divided into exonic-, intronic- and exon-intron-lncRNAs (Figure 5b). Intergenic lncRNAs were classified based on relative distance from the closest annotated PCG and assigned to one of three bins: ≤ 2 kb, ≤ 10 kb, or > 10 kb from the gene. We also annotated antisense lncRNAs that overlapped gene models based on intron-exon splice junctions derived from the genome-guided transcriptome reconstruction (see Methods; Additional file 14: Dataset S13). Since polyadenylated lncRNAs share biochemical features with coding mRNAs [58], we evaluated their exon number, transcript length, mean expression and spatiotemporal expression profiles. Sixty-seven percent of lncRNAs were unspliced (single exon) and these had a median length of 387 bp and showed a range of expression, including some that were highly expressed (Additional file 1: Figure S13). Both genic and intergenic lncRNAs showed strong enrichment of MNase HS at TSSs and TTSs, similar to PCGs (Figure 5c) and consistent with what has been shown in mammalian systems [59].

To test whether any of these early inflorescence-expressed lncRNAs were associated with genomic enhancer-like features, we leveraged a published set of ~1,500 candidate enhancer loci from maize leaf (V5) and husk tissues that were identified based on combined enrichment of histone 3 lysine 9 acetylation (H3K9ac) marks, high chromatin accessibility and low DNA methylation [16]. We expected a high degree of tissue specificity for the enhancer loci as well as the lncRNAs, but 45 inflorescence-expressed lncRNA loci overlapped with these enhancer regions (Additional file 12: Dataset S11). Expression profiles of these enhancer-associated lncRNAs were dynamic across tassel and ear development, with several accumulating in specific spatiotemporal patterns (Figure 5d). To investigate context-specific DNA methylation patterns at these loci, we re-analyzed and integrated publicly available genome-wide bisulfite sequencing data from 5 mm ear primordia [60]. While we observed a decrease in symmetric CG and CHG methylation from the proximal promoter into the gene body, there was a notably strong enrichment of asymmetric CHH methylation immediately upstream of the enhancer-associated lncRNA TSSs (Figure 5e). This asymmetric CHH methylation profile, known as an mCHH island, has been also shown to flank active PCGs as well as CNSs in plants, and has been hypothesized to mark the transition between heterochromatin to euchromatin [60]. Moreover, mCHH islands located upstream of TSSs were shown to be positively correlated with differential gene expression and tended to localize within open chromatin [61].

To explore potential *cis*-regulatory relationships between inflorescence-expressed lncRNAs and nearby PCGs, we analyzed correlation of their expression profiles across the eight spatiotemporal stages of immature tassel and ear development. Expression trajectories of PCGs were aligned with those of lncRNAs that were classified either as intronic or intergenic, and we prioritized lncRNA-PCG regulatory pairs based on congruence of expression patterns and their proximity to one another in the genome (Additional file 12: Dataset S11). We identified 78 lncRNA-PCG pairs that were i) positioned within a median distance of approximately 7 kb of each other and ii) were co-expressed across early maize inflorescence development (correlation coefficient > |0.8|). Among the PCGs that co-expressed with a lncRNA positioned within 2 kb, several encoded TFs including *wuschel-related homeobox 4 (wox4;* GRMZM6G260565) and other homeobox family members, as well as hormone synthesis and signaling components, e.g., *ethylene insensitive 2 (ein2;* GRMZM2G102754), *gibberellin 20-oxidase 2 (ga20ox2;* GRMZM2G127232) and *nine-cis-epoxycarotenoid dioxygenase 3(nced3;* GRMZM5G858784).

Based on the assumption that transcripts with similar expression trajectories might act in common pathways, we determined relationships between co-expressed lncRNA-PCG pairs (modules) based on hierarchical clustering (Additional file 1: Figure S15). For example, one module consisted of seven lncRNA-PCG pairs, each within 2 kb (Figure 5e). PCGs and lncRNAs in this module showed highest expression in 1 mm ear and 3-4 mm tassel primordia, consistent with a shift from indeterminate to determinate fate (Figure 5f) [32]. PCGs in this module included a SQUAMOSA BINDING PROTEIN (SBP) family TF *(sbp3;* GRMZM2G101499) and *ga20ox2.*

### Accessible chromatin-guided GWAS identifies novel SNP-trait associations for inflorescence architecture traits

Previous work in maize seedlings showed that while only 1–2 percent of the genome was MNase HS, this fraction accounted for more than 40 percent of the heritable variation [17]. We therefore hypothesized that our tassel and ear chromatin accessibility maps could help prioritize and/or assign biological function to trait-associated SNPs resulting from GWAS for inflorescence traits. We also speculated that using only those SNPs associated with regions of chromatin accessibility in a GWAS model could potentially help resolve novel SNP-trait associations, including smaller effect loci that may drive more subtle phenotypic variation. To investigate this, we first performed a GWAS analysis for two inflorescence architecture traits, tassel branch number (TBN) and ear row number (ERN), using publicly available phenotype data from the Nested Association Mapping (NAM) population and HapMap version (v) 3 SNP markers [62]. We also performed the GWAS analysis for these two traits using only SNPs that fell within accessible chromatin from the tassel and ear datasets combined. This latter analysis reduced the input SNP set to 15 million (approximately 19 percent of HapMap v3 SNPs).

For each set of markers, i) whole genome v3 SNPs and ii) SNPs within accessible chromatin regions, a stepwise model selection procedure was used [63]. Using the entire HapMap v3 SNP set, our GWAS model yielded 57 and 47 trait-associated SNPs for TBN and ERN, respectively. Interestingly, we identified different sets of associated SNPs for each trait from the two analyses. Using only SNPs within MNase HS regions identified a higher number of trait associations for TBN (n = 72) and ERN (n = 49) (Figure 6a). In addition, QTL identified using the reduced, MNase-based SNP set were generally positioned closer to known PCGs; i.e., the median distance from a trait-associated SNP to the nearest gene model was 649 and 2,273 bp for TBN and ERN, respectively, compared to 4.8 and 6.7 kb when using the entire v3 SNP set (Additional file 15: Dataset S14; Additional file 16: Dataset S15). This result was not surprising given that MNase HS regions are largely proximal to genes, however, it suggests that a reduced SNP set guided by accessible chromatin data could resolve novel associations that contribute to a phenotypic trait, which may not survive a more stringent multiple testing correction when a larger SNP set is used.

**Figure 6.**
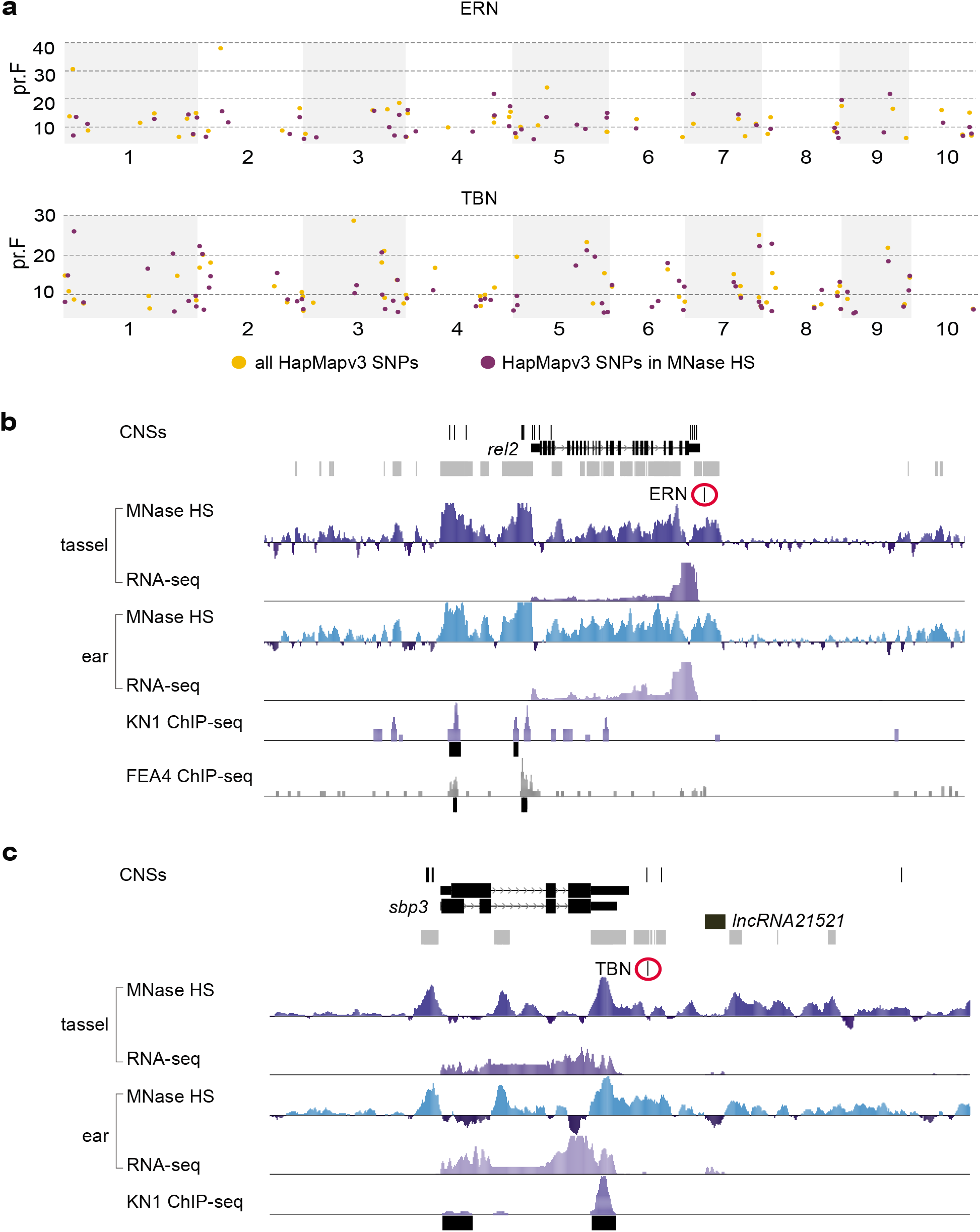
Novel trait-SNP associations for inflorescence architecture traits revealed using MNase-seq guided GWAS. Results from GWAS for inflorescence architecture traits, Ear Row Number (ERN) and Tassel Branch Number (TBN), comparing two sets of SNP markers, all HapMap v3 SNPs and the subset residing in MNase HS regions, revealed common and novel trait-SNP associations with candidate genes. **(a)** Significant SNP associations for ERN and TBN that were included in stepwise GWAS models using both marker sets displayed as a Z-browse view across the genome. Novel marker associations were found using the SNPs within MNase HS only for each trait. These include **(b)** *ramosa enhancer locus 2 (rel2),* which was adjacent to a SNP associated with ERN and **(c)** *sbp3,* an SBP TF in immediate proximity of a SNP association for TBN and just up-stream of co-expressed lncRNA21521. The grey track beneath the GWAS SNPs defines the MNase HS portion of the genome (tassel and ear combined), which was used to subset the SNP set.

There were nine trait-associated SNPs found in common between the two GWAS analyses for TBN, suggesting these are likely high-confidence associations that may contribute to phenotypic variation in tassel branching. One of these was located in the first large intron of GRMZM2G004690, an ortholog of the *ULTRAPETALA1* gene from Arabidopsis implicated in floral meristem determinacy [64] (Additional file 15: Dataset S14). In other cases, the two analyses identified unique SNPs that associated with the same protein-coding locus. For example, both analyses identified SNPs associated with ERN in the second to last intron of GRMZM2G049269, a gene annotated as a methyltransferase; this gene immediately flanks *Gibberellic acid (GA) 20-oxidase 5* (GRMZM2G049418), which we showed to be differentially regulated between tassel and ear (Figure 2d). SNP-trait associations with key developmental genes implicated in maize inflorescence architecture were identified with the MNase HS-guided GWAS analysis, but not when using the entire v3 SNP set. For example, *barren inflorescence 1* (bif1), a classical maize gene that plays a major role in tassel branching [65], harbored a SNP association for TBN. Another gene that regulates inflorescence branching in maize, *ramosa enhancer locus 2 (rel2;* GRMZM2G042992) [66], was identified as associated with a SNP for ERN (Figure 6b). A SNP-trait association for TBN was identified at *sbp3* (GRMZM2G101499) using only MNase SNPs (Figure 6c). This SBP TF was co-expressed with lncRNA21521 positioned immediately downstream, and both were part of a co-expression module associated with meristem determinacy (Figure 5e).

Since this analysis was performed using the NAM population, we expect that the haplotype blocks will limit our ability to resolve causal SNPs, regardless of subsetting the functional genome. We also tested this approach in a small association panel, the 282 line Goodman-Buckler maize diversity panel [67], using GWAS results for TBN from [68]. For the subset of markers present in tassel MNase HS regions (n = 71,024), new False Discovery Rate (FDR)-adjusted P-values were calculated using the Benjamini and Hochberg [69] procedure. Overall, markers within tassel MNase regions showed a greater reduction in adjusted p-values compared to the genome-wide analysis (Additional File 1: Figure S16). Since a decrease in FDR-adjusted p-values could be due to the reduced number of marker association tests, we calculated new FDR adjusted p-values for 1,000 random samples of 71,024 markers. For each replicate, we calculated a statistic we denote as lambda (λ; see Methods and Additional File 1: Figure S16 for equation), which is a ratio of the FDR adjusted p-value for a given marker when a reduced marker set is used, compared to the full marker set. Overall, the markers in tassel MNase regions had greater reductions in adjusted *p*-values compared to random samples (four standard deviations from the mean of random iterations; Additional File 1: Figure S16). This result suggests that markers present in early tassel MNase HS regions could be of higher biological relevance for phenotypic variation in TBN than those scattered randomly across the genome. It also demonstrates how the reduction in total number of markers tested can facilitate identification of less significant, but potentially biologically interesting, associations. This approach for reducing markers could improve the efficiency for marker assisted selection.

## Discussion

Precise developmental transitions, such as those underlying distinct branching morphologies of maize tassel and ear, are regulated through dynamic interactions between TFs and non-coding elements of the genome. Regulatory variation across spatial and temporal scales underpins morphological diversity in evolution, and can be harnessed to achieve optimized phenotypic outputs in crops through precision breeding and/or engineering. In maize, several core regulatory factors that control inflorescence patterning have been identified through classical mutagenesis studies. However, most of these genes have not been associated with natural variation in inflorescence architecture, despite the high heritability of these traits in large association panels [70]. Since perturbations in these core regulators typically result in extreme and/or pleiotropic phenotypes [33,71], it is likely that natural variation in cis-regulatory sequences modulate these genes and/or their targets. A major challenge in genomics-enabled crop improvement is functional annotation of *cis*-regulatory elements in crop genomes, and the ability to harness these sequences to fine-tune specific pathways with little perturbation to the complex networks within which they reside [72]. These genome-wide analyses of chromatin accessibility and nucleosome occupancy, tissue-specific regulatory RNAs and TF-DNA predictions provide a foundation for exploring the functional maize genome that underlies early developmental decisions in tassel and ear organogenesis.

Maize is unique among the major cereal crops in having separate male and female structures, which has facilitated hybrid seed production. Over the past century, maize improvement has gone hand-in-hand with selection of smaller tassels that take up fewer resources and less real estate (in terms of light interception), and larger, more productive ears [73]. Since the tassel and ear develop by way of a common developmental program, further improvement of ear traits by advanced breeding and engineering will require decoupling of this program, for example, via tassel- or ear-specific promoters or regulatory elements. Therefore, understanding how the same genes are regulated differently in tassel and ear, and harnessing this specificity to control one over the other, will advance breeding efforts in maize. In addition to agronomic applications, the maize tassel and ear represent an ideal system for studying the control of meristem determinacy and regulation of axillary branching in plant development. The core developmental program that underlies the two maize inflorescences is also shared with those of other grasses; the same meristem types progress in sequential order, but with variation on determinacy and fate. For example, the indeterminate branch meristems (BMs) that grow out into long branches at the base of the tassel, are determinate in the ear.

Our analysis of differentially regulated genes between tassel and ear was consistent with a determinacy program imposed on BMs in the ear, allowing for kernels born on short branches in organized rows. We have some understanding of the key players that suppress branching in the inflorescence such as the *ramosa (ra)* genes [32,71] and tb1, however little is known about the mechanisms or what other factors they interact with. A number of other transcriptional regulators were preferentially expressed in the ear during early development, and there was tissue-specific chromatin accessibility associated with them. Among the most differentially regulated genes were five TCP TFs, including *tb1* and four others that have not yet been functionally characterized. We also found that TCP binding motifs were enriched in promoters that were more accessible in ear. TCPs are among the bHLH (basic Helix-Loop-Helix) family of TFs, and are further grouped into class I or class II TCPs. Aside from tb1, the other four TCPs that were differentially regulated in ear are class I.

There are several examples of class II TCPs regulating plant form in crops; suppression of axillary meristem or tiller outgrowth in grasses by *tb1* [27,74], lateral branch angle in maize tassels by *wavy auricle on blade 1/ branch angle defective 1 (wab1/ bad1)* [75,76], and floral symmetry in sunflower by *HaCYC2* [77]. Recently, it was shown that a class II TCP, MULTISEEDED 1 (MSD1), regulates floral fertility and therefore grain number in sorghum, by mediating accumulation of jasmonic acid in the panicle [78]. Class I TCPs have been largely implicated in both abiotic and biotic stress responses [79,80], and there is evidence emerging for their role in hormone pathways that intersect developmental processes. In Arabidopsis, TCP14 and 15 are class I TCPs that physically interact with SPINDLY (SPY) to activate cytokinin response genes in leaves and flowers [81], and TCP20 binds and regulates *LIPOXYGENASE 2 (LOX2),* which encodes a key enzyme in JA biosynthesis [82]. In our findings, genes implicated in JA biosynthesis were overrepresented among those up-regulated in tassel compared to ear. TCPs can form homo- and hetero-dimers to achieve different binding specificities to target genes [83,84]. Class I TCPs 20, 6 and 11, predicted to be functionally redundant, bind a conserved sequence GCCCR [85], which closely matches the *de novo* PWM enriched in ear accessible footprints (RGCCS; Additional file 1: Table S5). TCP20 also directly binds the promoter of TCP9, another class I TF [82]. We observed multiple occurrences of TCP binding sites in the promoters of class I TCPs preferentially transcribed in ear, indicating several layers of control by TCP family members.

Of the five TCP TFs that we identified as being differentially regulated in the ear, only the well-characterized *tb1* gene was associated with any functional data in maize. This highlights the utility of genomics-enabled research for gene discovery, particularly when viewed in a context-specific manner. There are approximately 2,700 TFs in the maize genome and we only have experimental evidence on how a small fraction of them function [3,86]. Context-specific gene regulatory networks (co-expression and TF-DNA binding data) can help in predicting functions of uncharacterized TFs and other genes of unknown function based on their position in the network [18,87]. Genome-wide TF-DNA binding data have been limited in plants, especially in crop species, largely due to inherent difficulties in performing *in planta* methods such as ChIP-seq [88]. We showed here that there is extensive overlap of *in planta* TF binding data with MNase HS in the same tissue type, suggesting that accessible regions can be viewed as a proxy for TF binding. We also observed increased accessibility in promoters of genes bound by two TFs based on experimental data, indicating that degree of accessibility scaled with number of binding factors. Therefore, these tissue-specific accessibility maps can help drive computational-based predictions of TF binding and refine results from *in vitro* methods such as DNA Affinity and Purification sequencing (DAP-seq) [9,89]. Along with co-expression networks of genes from the same tissue type, the accessible chromatin maps provide a basis for strengthening TF-DNA interaction predictions.

There is increasing evidence for lncRNA-mediated regulation of genome structure in both plants and animals, e.g., through nucleosome positioning, interaction with chromatin remodelers and chromatin looping [53,55,90]. In animal systems, it has been demonstrated that lncRNAs transcribed from enhancer loci, called enhancer RNAs (eRNAs), can modulate the expression of target genes [50,91]. While regulation of PCGs by lncRNAs can occur in both *cis* and *trans,* several studies have highlighted control of the local regulatory “neighborhood” by lncRNAs [91–93]. Various mechanisms by which lncRNAs regulate expression of proximal PCGs have been reported [94], as well as transcriptional regulation by the same ‘divergent’ promoter [95,96]. In Arabidopsis, expression of an intergenic lncRNA locus, *APOLO,* is auxin-dependent and destabilizes a chromatin loop to the promoter of *PINOID (PID),* a key regulator of polar auxin transport [53], allowing its expression.

LncRNAs in both plants and animals tend to share little evolutionary origin, biological function, or molecular mechanism [58]. Therefore, context specificity is critical to elucidating the function of these non-coding elements in growth and development. The set of 2,679 lncRNAs annotated for early stages of developing inflorescence primordia extend lncRNA catalogs described from other maize tissues [97,98]. LncRNAs have also been implicated in fine-tuning homeostasis and developmental programs. For example, in embryonic stem cells, massive coordinated changes in expression of proximal lncRNAs and PCGs off of divergent promoters underpinned organ differentiation programs [96] and in mouse, lncRNAs flanking HOX gene clusters were shown to modulate specificity of HOX gene expression during development [99]. In our analyses, we observed modules of co-expressed PCG-lncRNA pairs, including one module that exhibited an expression profile that was coordinated with spikelet pair meristem determinacy [32]. Within this module, an SBP TF, *sbp3,* was co-expressed with a lncRNA immediately downstream. SBP TFs play key roles in developmental processes, however *sbp3* has not been functionally characterized. Further, its association with a GWAS SNP for TBN and its enhanced expression in ear, suggest that *sbp3* and the co-expressed lncRNA are potential candidates for regulating branching (Figure 6c). A genome-wide study for associations with complex traits in humans uncovered numerous lncRNAs, which were often associated with control of adjacent PCGs [100]. Further experimental validation is necessary to link specific gene regulation to proximal lncRNAs in maize inflorescence development.

The ENCODE project revealed that much variation attributable to major human disease is associated with accessible regions in the human genome [22]. Since then, work in maize showed that MNase HS regions explained 40% of the heritable variation in complex traits [17] and in Arabidopsis, DNase I HS regions were enriched for GWAS hits [18]. Our rationale for performing GWAS using only markers within MNase accessible DNA was to test whether we could recover variants that fell within functional regions of the genome, and potentially even pinpoint causal variants that would otherwise be removed from the stepwise selection model in favor of a different SNP in linkage disequilibrium, or fall below the significance threshold of multiple testing with large numbers of markers. Furthermore, the MNase HS regions used reflect chromatin accessibility in the tissue type affected by the inflorescence traits of interest. By limiting HapMapv3 SNPs to MNase HS regions, we used only 19% of the markers. The resulting SNP-trait associations that were uncovered using the reduced marker set were different from our GWAS results using the entire set of 83 million markers. Since a stepwise model selection procedure was used, we would not expect the genome-wide model to include all SNPs from the MNase-guided model. Among trait-associated SNPs in the reduced set GWAS model were several that resided either within or proximal to known developmental genes that have been implicated in inflorescence architecture. One of these was *rel2,* a gene that encodes a TOPLESS-like co-repressor that was identified in a modifier screen as an enhancer of *ramosa 1* (enhanced branching [66]). Notably, here *rel2* associated with ERN, which is consistent with recent work that showed mutants in this gene also display a fasciated ear phenotype, which is developmentally linked to ERN [101]. Experimental validation is needed in order to test whether the other trait-SNP associations identified by the MNase-seq driven GWAS analysis underlie variation in maize inflorescence morphology. We hypothesize that with a large association panel, reducing the marker set to those in functional regions could improve resolution of genomic selection models by i) prioritizing functional SNPs, ii) lowering the significance threshold required to declare a marker significant, and iii) reducing the computational load.

In summary, the functional maps described here provide a foundation for pinpointing key regulatory signatures that underlie growth, development, and morphological diversity in reproductive structures of maize, the most important cereal crop. The MNase-seq profiles offer genome-wide views of the regulatory space, including both TF binding sites as well as nucleosome occupancy, which can be leveraged for making informed predictions on functional DNA, and as a complement to many other genomics analyses. One timely application is guiding gene editing strategies by determining accessibility of target regions. Our integrated analyses of gene regulation during early inflorescence organogenesis defined targets for manipulating architecture, and provided insights into tassel- and ear-specific developmental programs that can be harnessed for crop improvement.

## Conclusions

Our ability to predict and achieve desired crop phenotypes through alteration of genetic elements is enhanced by context-specific regulatory maps targeted to traits of interest. The systems-level analyses presented here explore the *cis*-regulatory landscape underlying developmental transitions that occur during differentiation of male and female inflorecences of maize. Chromatin accessibility and nucleosome occupancy maps in these young, meristematic tissues revealed regulatory signatures specific to tassel and ear development including differentially regulated members of certain TF families, and predicted TF binding sites to footprint level resolution. On a genome-wide scale, differences in accessibility were observed with tassel-specific HS signatures enriched in intergenic regions while ear-specific HS were more prominent in gene body features. Several intergenic accessible sites coincided with predicted promoters of spatiotemporally expressed lncRNAs, combinatorial TF binding, and trait-SNP associations from GWAS, prioritizing them as potential long-range regulatory elements. These integrated functional maps and associated analyses can be directly leveraged for targeted crop improvement in maize and are readily translatable to other economically important cereal crops.

## Methods

### Plant material

Maize (Zea *mays)* B73 plants were grown in a greenhouse environment (27°C/23°C day/night) at the DDPSC Integrated Plant Growth Facility. Three seedlings per 2 gallon pot (in Pro-Mix BRK-20 soil) were harvested 4-5 weeks and 6-7 weeks after sowing to collect tassel and ear primordia, respectively. For each tissue type, approximately 170 inflorescence primordia (between 1-5mm in size) were hand-dissected and pooled into each of two biological replicates. Primordia were immediately flash frozen in liquid nitrogen, and stored at −80°C.

### Nuclei isolation

Nuclei were isolated as previously described [17]. Briefly, one gram of tissue was ground in liquid nitrogen with a mortar and pestle. Cross-linking was performed using 10 ml ice-cold fixation buffer (15 mM Pipes·NaOH at pH 6.8, 0.32 M sorbitol, 80 mM KCl, 20 mM NaCl, 0.5 mM EGTA, 2mM EDTA, 1 mM DTT, 0.15 mM spermine, and 0.5 mM spermidine) containing 1% formaldehyde (10 min incubation with rotation) at room temperature and fixation was stopped using 0.5 mL of 2.5 M glycine. Triton X-100, a non-ionic surfactant, was added and the tube rotated for 10 min to release nuclei from the cells. The μsuspension was filtered through one layer of Miracloth, placed in a 15 ml falcon tube, and then split into two tubes. A percoll gradient separation using 50% (vol/vol) percoll in phosphate buffer saline (PBS) followed by low speed centrifugation at 3,000 × g for 15 min at 4°C was used to enrich for nuclei. Using a serological pipette extended to the bottom of the tube, 4 mL of 50% (vol/vol) percoll was slowly added underneath the filtered nuclei suspension in PBS. Phase separation of nuclei was performed by centrifugation at 3,000 × g for 15 min at 4°C, and nuclei at the percoll-PBS interface were transferred to a 15 ml falcon tube. PBS was added to dilute the nuclei suspension to a final volume of 15 ml. Nuclei were then pelleted by centrifugation at 2,000 × g for 10 min at 4°C, and pellets resuspended in 2.1 mL MNase digestion buffer (MDB, sterilized with 0.2 μm filter, 50 mM HEPES-NaOH pH 7.6, 12.5% glycerol, 25 mM KCl, 4 mM MgCl2, 1 mM CaCl2). Nuclei were flash frozen in liquid nitrogen and stored at −80°C until use. To check nuclei quality, the remaining 100 μl were stained with DAPI and imaged as follows: 8 μL of DAPI-stained nuclei in MDB were mounted on a glass slide in VectaShield+DAPI medium and subjected to 3D deconvolution imaging using a DeltaVision (GE Healthcare) equipped with a 60X lens. Images of nuclei preps were produced using maximum intensity projections of the 3D datasets.

### MNase digestion, DNA extraction, and library preparation and sequencing

Intact nuclei were digested with a range of MNase concentrations in order to determine the optimal enzyme concentration for heavy and light digests, as described by [34]. After MNase titration, nuclei were digested by adding MNase to 30 U/ml (light) or 7290 U/ml (heavy), and incubated for 5 min at room temperature. Digestion reactions were terminated by adding 10 mM EGTA. Nuclei were reverse crosslinked with 1% SDS and 100 mg/mL proteinase K by incubation overnight at 65°C. Then, DNA was extracted using 25:24:1 phenol-chloroform-isoamylalcohol, ethanol precipitation and finally resuspended in TER (TE with 40 ug/mL RNase A). Mononucleosome-sized DNA fragments (~150 bp) and smaller were size-selected using AMPure XP beads by adding 90% sample volume of beads (Additional file 1: Figure S1). Size-selected DNA fragments were used to prepare eight MNase sequencing libraries with the NEBNext Ultra DNA Library Prep Kit for Illumina (NEB), following manufacturer’s instructions (2 biological replicates x 2 MNase concentrations x 2 tissues). For tassel and ear, the four libraries were barcoded and pooled into four lanes of sequencing on the HiSeq2500 at the FSU sequencing facility.

### MNase-seq data mapping and analyses

Paired-end reads were trimmed and mapped to the maize B73 AGPv3 reference genome as described in the MNase-seq processing pipeline [34] (Additional File 1: Table S1). Read coverage (in reads per million (RPM)) was calculated in 25 bp windows as described in [34] for input into bedtools. DNS was calculated as the difference of mean normalized depth (RPM) between the light and heavy digests. Positive values correspond to accessible chromatin and negative values correspond to closed regions. Read coverages were highly correlated between biological replicates for both light (r > 0.9) and heavy (r > 0.73) digested libraries. The read coverage for the MNase datasets (DNS and LCS) calculated in sliding windows (window size = 1 Mb and step size = 100 kb) was used to visualize accessible regions across the genome using *Circos* [102]. Positive and negative values were plotted for DNS and positive Z-scores were plotted for the LCS data.

MNase HS regions were identified in each of the biological replicates and combined data sets using the genomic segmentation algorithm, iSeg [38] and output files were processed using custom scripts. Our analysis applied a range of biological cutoff (bc) stringencies in calling MNase HS or resistant regions; from bc 0.5 (lowest stringency) to bc 3.0 (highest stringency) (Additional file 1: Tables S2 and S3; Additional files 2 and 3: Datasets S1 and S2). To validate results from iSeg, we also performed MACS2 [103] using the light digest as “treatment” and heavy digest as “control”, and parameters “-g 2.3e9 -nolamda -q 0.01”. Using a bc of 2.0 on DNS data, iSeg identified > 90% of the HS regions called by MACS2, in addition to many more putative regulatory regions (Additional file 1: Tables S2 and S3). Comparing these accessible regions with ChIP-seq data for FEA4, 85% of high-confidence FEA4 binding sites overlapped with accessible regions called by iSeg in tassel compared to 42% called by MACS2 (Additional file 1: Figure S2). Decreasing the iSeg peak-calling stringency from bc 3.0 to 2.0 increased the overlap with TF-bound regions, but did not significantly increase noise (Additional file 1: Figure S2). Therefore, bc 2.0 was chosen as the default cut-off used for analyses in the manuscript.

At bc 2.0, ~77% of DNS peaks overlapped between biological replicates and high-confidence HS regions were called if their midpoints were within 300 bp of each other (Additional file 4: Dataset S3). Manual inspection in a genome browser revealed several cases where MNase HS was visible in the two biological replicates, but only called as a peak in one likely due to coverage threshold (Additional file 1: Figure S3). To enhance depth of coverage for resolution of dynamic accessibility and footprints of bound TFs, we also combined reads from the two replicates and iSeg was used to call MNase HS on the combined datasets as well as for individual replicates. The MNase-seq data were also mapped to the maize RefGen_v5 genome, and iSeg coordinates for both DNS and LCS are available in GEO (Additional File 5: Dataset S4).

To map distributions of MNase HS sites across genomic features, we used the Bioconductor packages *IRanges, GenomicRanges,* and *GenomicFeatures* [104]. A transcripts database object was generated based on the AGPv3.31 reference gene annotation using the function *makeTxDbFromGFF,* and coordinates of genomic features were determined corresponding to PCG models. For each gene model, only primary transcripts were used and duplicated 5’ UTRs were removed. The total length of each genomic region (in bp) was calculated using the function *sum.* The midpoint of the MNase HS sites were computed in R, converted to a *GRanges* object, and intersected with respective genomic regions using the function *findOverlaps* with the options *type=within, ignore.strand=F*. Intergenic MNase HS sites falling within the reference repeats annotation file (AGPv3.31) were excluded from the analysis.

### Transposable element content

The maize TE annotation file (AGPv4) was downloaded from Gramene release 61 [41]. TEs annotated on chromosomes 1 to 10 were selected and converted to AGPv3 using the tool CrossMap v.0.3.7 [105] and the chain file from Gramene release 61. The resulting TE coordinates projected to AGPv3 scaffolds were removed and not considered in the downstream analysis. TE coordinates with unassigned strand “*” were considered as located on the sense strand. Intergenic TEs were selected, calssified and intersected with the midpoints of ear and tassel MNase HS regions (DNS) using the R package *GenomicRanges* (options: type = ‘within’, ignore.strand = T). The enrichment analysis was conducted using the Bioconductor package *regioneR* [106] based on permutation tests (n = 1,000). A randomization strategy to compare intergenic genomic regions was performed using the function circularRandomizeRegions.

### RNA-seq data analysis

Publicly available RNA-seq data from developing maize tassel and ear primordia were downloaded from the NCBI Sequence Read Archive (SRA) database, BioProject PRJNA219741 (Run IDs: SRR999054, SRR999053, SRR999052, SRR999045, SRR999044, SRR999043, SRR999042, SRR999041, SRR999040, SRR999039, SRR999038, SRR999037, SRR999036, SRR999035, SRR999034, SRR999033, SRR999032) [32] and re-analyzed. We used ~500 M high-quality reads from these libraries to reconstruct the maize transcriptome using a genome-guided approach with HISAT2 v2.0.6 [107] and StringTie v1.2.3 [108]. Two mapping iteration steps were performed on each library: a first round of mapping to the reference B73 maize genome (AGPv3.31) from Ensembl Genomes [109] with options ‘--*novel-splicesite-outfile, --dta’* and without a reference gene annotation file. Based on information from this first alignment, a database of unique splice junctions was created to maximize the number of mapped reads in the second mapping iteration using the option ‘*--novel-splicesite-infile’*. Then for each library we used the reference gene models (AGPv3.31) to guide the transcriptome reconstruction (assembly) and merged (using option --merge) all assemblies to create a non-redundant reference for this dataset. The transcript sequences (FASTA) and the gene models (GFF) were extracted using the *gffread* utility distributed with the program Cufflinks [110].

MNase HS and expression data for each PCG were correlated using the mean normalized depth (reads per million) for the heavy and light MNase digests and the StringTie v1.3.5 TPM values for 1-2 mm tassel and ear data. Using the R package *travis* (https://github.com/dvera/travis), the normalized coverage (RPM) was calculated for light digests, heavy digest and the differential (light-heavy) at the midpoint for 25 bp genomic windows.

### Differential DNS analysis

Differential accessibility within 500 bp upstream of the TSS of PCGs was calculated between tassel and ear was by computing the MNase HS value (tassel DHS – ear DHS) in 25 bp windows. Based on these values, changes of > 1 or < −1 were defined as differentially accessible in tassel or ear, respectively. Genes characterized by differential accessibility in their promoter were filtered to include those with >100 read coverage selected based on the RNA-seq data. A fold change of |1.5| between the two tissues was used to exclude genes with minimal expression changes [111].

### Analysis of regulatory elements within MNase HS regions

ChIP-seq data for FEA4 and KN1 are publicly available in the NCBI Gene Expression Omnibus (accession numbers GSE61954 and GSE39161, respectively). Genome-wide binding sites were determined for FEA4 using MACS v2 [103] with parameters ‘-f BAM -g 2.3e9 -B -m 10 30 -p 1.00e-5’. High-confidence peaks were called if peak summits from two biological replicates were within 300 bp. For KN1, peak coordinates were lifted to maize AGPv3 using the ‘liftOver’ function in the UCSC genome browser. High-confidence peaks required a minimum overlap of 50 percent between two biological replicates. Regions were designated as co-bound by FEA4 and KN1 if midpoints of high-confidence peaks were within 500 bp.

The MEME suite v5.0.5 (www.meme-suite.org/; [112]) was used for motif analyses. The consensus binding site for FEA4 was determined using MEME-ChIP with default settings to mine genomic regions 200 bp around summits of high-confidence FEA4 ChIP-seq peaks (n = 2,163). Random 200 bp maize genomic DNA fragments were used as background. The resulting FEA4 PWM was located within FEA4 ChIP-seq peaks using Find Individual Motif Occurrences (FIMO; threshold = 0.0005) in MEME. The custom R package *genmat* (https://github.com/dvera/genmat) was used to generate density plots of MNase HS around FEA4 binding. HS regions overlapping FEA4 binding sites called by iSeg (bc 2.0) in tassel and ear were used to scan for enriched proximal motifs using two *de novo* motif finding tools: MEME-ChIP and Hypergeometric Optimization of Motif EnRichment (HOMER; http://homer.ucsd.edu/). MEME-ChIP was used with settings ‘-ccut 0’ and inputting a random set of genomic sequences as a control. A parallel analysis used a random set of iSeg-called maize sequences as the input compared to random genomic controls. Input and control sequences were repeat-masked. The Tomtom function in MEME was used to assign resulting motifs to best matches in the JASPAR core plant PWMs (2018; http://jaspar.genereg.net). We used the findmotifsgenome.pl function in HOMER with setting ‘-size given’ to report best matches to known plant motifs within the HOMER database. Resulting PWMs from HOMER were also matched to the JASPAR core plant PWMs.

To resolve locations of putative TF footprints we mapped the midpoints of small (<131 bp) fragments reads generated from light MNase digests. To find the fragment center hot spots, we converted the single-base fragment center data using a sliding window average, where the total number of fragment centers in a 21 bp window was assigned to the central 5 bp in that 21 bp window, followed by repeating a 5 bp step size. To aid visualization of the fragment center populations on a genome browser, this procedure was also performed for the 5’ and 3’ ends of the light digest MNase fragments (0-130 bp). The resulting fragment center values showed local peaks that were segmented using the iSeg peak-calling program. *De-novo* motif discovery using these footprints was performed using Discriminative Regular Expression Motif Elicitation (DREME) in the MEME suite using default settings. Random background sequences of equal size from the same promoters were used as control. Tomtom was used to find the best matches to motifs in the JASPAR plant database. The resulting PWMs were located within footprints using FIMO (threshold = 0.005).

### Annotation and quantification of long non-coding RNAs

High-confidence lncRNAs were annotated according to De Quattro et al. [56,113]. Non-redundant transcripts assembled from the RNA-seq data were scanned using published criteria for defining lncRNA transcripts [58,114]: we filtered out sequences that i) were less than 200 bp, ii) were open reading frame encoding proteins with more than 100 amino acids, iii) were known functional protein domains deposited in the database Pfam v30 [115] (BLASTX cutoff E-value ≤ 1.0E −3), iv) had the potential to encode functional proteins based on the program CPC [116], and/or v) had homology with maize structural RNA domains deposited in the database Rfam v12 [117] (BLASTN cutoff E-value 1.0E ≤ −10). 15,606 potential noncoding transcripts were discarded due to protein coding capacity. To exclude potential small RNA precursors from our analysis (e.g., microRNAs and phasiRNAs), transcripts passing these filters were further processed using publicly available small RNA datasets (> 37 M processed small RNA reads) from immature B73 tassel and ear (~0.5 – 2 cm) [118] (SRA BioProject PRJNA119855, run ID: SRR032087, SRR03209) and A619 tassel primordia (0.5 – 1 cm) [119] (SRA BioProject PRJNA230402, run ID: SRR1041542, SRR1041543, SRR1041544). Small RNA reads (18-34 nt in length) were mapped against the lncRNA sequences using PatMaN [120] and processed with custom scripts in R. LncRNA sequences hosting ≥ 20 different small RNA reads were discarded. To reduce noise from spuriously transcribed RNAs, we considered only transcripts with a sum expression cut-off equal to or greater than 10 TPM across the entire dataset. Ten percent of lncRNA prediction was supported by Sanger sequencing (> 80% alignment identity and 50% alignment coverage) using maize NCBI ESTs (n=393,418) (Additional file 13: Dataset S12).

High-confidence lncRNAs were classified based on their relative genomic location to protein coding genes. We defined overlaps between lncRNAs and PCGs using the Bioconductor package, *GenomicRanges.* LncRNAs were considered genic if they were located i) entirely within exons, ii) entirely within introns, or iii) spanning exon-intron boundaries. LncRNAs overlapping a PCG model in the antisense orientation were classified as antisense. Expression profiles of lncRNAs were determined by calculating TPM values across the eight stages of maize inflorescence development using Salmon v0.8.2 in mapping-based mode [121] and the sequences derived from the transcriptome reconstruction. Default parameters were used except the option --numBoostraps 30. Salmon output files were imported into R and processed using the Bioconductor package *tximport* [122]. Meristem-specific expression of lncRNAs was assessed by the Shannon Entropy (SH) metric using the Bioconductor package *BioQC* [123] and the TPM expression data.

To correlate expression levels of the lncRNAs with their closest PCGs, pairwise co-expression analyses were performed across each lncRNA – PCG pair independently for each of these four classes of lncRNAs: intronic-lncRNAs and intergenic lncRNAs < 2 kb, < 10 kb, and >10 kb. First, expression values of PCGs and lncRNAs were normalized by log transformation using the function *rlog* in the Bioconductor package *DESeq2* [124]. For each of the four lncRNA types, a matrix of lncRNA-PCG pairs was generated. For each pair, their expression profiles and correlation between them were determined using a biweight mid-correlation method with the function *bicor* from the Bioconductor package *WGCNA* [125]. The lncRNA-PCG pairs with a correlation coefficient >|0.8| were kept and hierarchically clustered to determine modules of co-expressed pairs.

### Analysis of whole genome bisulfìte sequencing data

Public raw data from whole-genome bisulfite sequencing (paired-ends; 100 bp) of B73 immature ears (~5 mm in length) were downloaded from the NCBI Sequence Read Archive using the run ID SRR2079442 [60]. Raw data were processed for quality using the program trim_galore (v.0.4.4_dev) with the parameters ‘-q 20 --length 75 --trim-n --retain_unpaired’. Quality reads were mapped to the maize AGPv3 reference genome using the program BSMAP [126] with the options ‘-r 2 -v 5’. DNA methylation level at single cytosine in the three methylation contexts (mCG, mCHG, mCHH) was extracted using the script methratio.py included in BSMAP.

### Functional enrichment analysis

Maize-GAMER [127] AGPv3 aggregate annotation file was downloaded from Cyverse (DOI: doi.org/10.7946/P2S62P) and used as input for the gene set enrichment analysis. The function *enricher* from the Bioconductor package *clusterProfiler* [128] was used with default parameters and Benjamini-Hochberg multiple test correction. Annotation of GO terms were retrieved from the database QuickGO (https://www.ebi.ac.uk/QuickGO/). Maize TF annotations were downloaded from Grassius [86] and PlantTFDB [129]. The two databases were merged and filtered for duplicated entries and used as input in the enrichment analysis by hypergeometric test as described earlier for the GO terms.

### Genome-wide association study for TBN and ERN

The dependent variable for each GWAS model was the residuals from a joint linkage analysis model fitted in [130] containing all quantitative trait loci associated with the trait of interest except for those on the given chromosome. Because the SNP markers were genotyped on only the NAM founders, we used a procedure similar to that described in [130–132] to project these markers onto the entire NAM population. For each set of markers, a stepwise model selection procedure implemented in the software TASSEL v5.0 [133] was used. Entry and exit criteria for each marker set was determined through a permutation procedure described in [134], where 100 permutations were conducted. To ensure the detection and quantification of both large- and moderate-effect genomic signals, the empirical entry/exit thresholds were determined at α = 0.20. Data were visualized in *Zbrowse,* an interactive GWAS viewer (www.baxterlab.org) [135].

To test the performance of MNase SNPs within the 282 Goodman-Buckler maize diversity panel, we used GWAS results from Rice et al. (2020; [68]) for TBN and calculated new False Discovery Rate (FDR)-adjusted p-values for SNPs within tassel MNase HS regions and 1,000 random subsets (n = 71,024) using the Benjamini and Hochberg procedure ([69]; alpha = 0.05). For each of the 1,000 replicate iterations, we calculated the following statistic, lambda (λ):

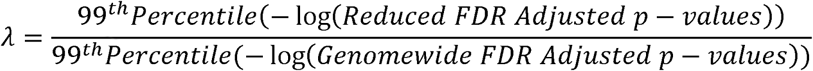

A λ of one means that there was no change in the 99^th^ percentile, i.e., the top 1% most significant SNPs, compared to the values in a genome-wide analysis. The higher the λ, the greater the FDR-adjusted *p*-value of the 99^th^ percentile decreased.

## Declarations

### Ethics approval and consent to participate

Not applicable.

### Consent for publication

Not applicable.

### Availability of data and materials

All sequence-based datasets generated from this study along with associated metadata and processed data, including map locations for HS regions in maize AGPv3 and RefGen_v5 and GWAS data, have been deposited to NCBI Gene Expression Omnibus (GEO) under the accession number GSE136685 [136]. Due to large file sizes, Additional files 2, 3, 4, 6, 11 and 16 have also been deposited in GEO at GSE136685 [136].

For the reference datasets used in the manuscript:

Rodgers-Melnick E. Open chromatin reveals the functional maize genome. Maize shoot MNase-seq. SRA302258. https://www.ncbi.nlm.nih.gov/sra/SRA302258 (2016) [17].

Eveland AL. Regulatory modules controlling maize inflorescence architecture. RNA-seq. GSE51050. https://www.ncbi.nlm.nih.gov/geo/query/acc.cgi?acc=GSE51050 (2014) [32].

Pautler M. FASCIATED EAR4 encodes a bZIP transcription factor that regulates shoot meristem size in maize. FEA4 ChIP-seq. GSE61954. https://www.ncbi.nlm.nih.gov/geo/query/acc.cgi?acc=GSE61954 (2015) [35].

Bolduc N. Unraveling the KNOTTED1 regulatory network in maize meristems. KN1 ChIP-seq. GSE39161. https://www.ncbi.nlm.nih.gov/geo/query/acc.cgi?acc=GSE39161 (2012) [36].

Li Q. RNA-directed DNA methylation enforces boundaries between heterochromatin and euchromatin in the maize genome. Bisulfite-seq. SRR2079442. https://www.ncbi.nlm.nih.gov/sra/?term=SRR2079442 (2017) [60].

### Authors’ contributions

HWB and ALE designed the research. RKP, EB, DV, MS, P-YL, and BRR performed data analyses. HWB, JZ, AEL and ALE advised data analyses. MS performed the MNase-seq experiments with advice from DV and HWB. EB annotated the lncRNAs. BRR, PB, AEL performed and advised the GWAS. RKP, EB, and ALE wrote the paper. All authors contributed to editing the final manuscript.

## Acknowledgements

The authors would like to thank Noah Fahlgren and the bioinformatics core at DDPSC, the Center for Genomics and Personalized Medicine at FSU, and the National Center for Supercomputing Applications at UIUC for computational support. Also, special thanks to Kevin Reilly and the Integrated Growth Facility at the DDPSC for plant care.

## Additional files

**file names starting with GSE136685 are available at NCBI GEO

**Additional file 1.** Supplemental figures (S1-S16) and tables (S1-S6). (**Additional_file1.pdf**)

**Additional file 2: Dataset S1**. Coordinates of all MNase HS peaks (including analyses from DNS and LCS; AGPv3; iSeg bc 2.0). (**GSE136685_AdditionalFile2_DatasetS1.txt**)

**Additional file 3: Dataset S2**. Coordinates of all MNase HS peaks (including analyses from DNS and LCS; AGPv3; iSeg bc 3.0). (**GSE136685_AdditionalFile3_DatasetS2.txt**)

**Additional file 4: Dataset S3**. High confidence MNase HS peaks (including analyses from DNS and LCS; AGPv3; iSeg bc 2.0). (**GSE136685_AdditionalFile4_DatasetS3.txt**)

**Additional file 5: Dataset S4**. Coordinates of all MNase HS peaks (including analyses from DNS and LCS; iSeg bc 2.0) mapped to the maize reference genome RefGen_v5. (**GSE136685_AdditionalFile5_DatasetS4.txt**)

**Additional file 6: Dataset S5**. Unique MNase HS peaks (DNS) in tassel and ear primordia (iSeg bc 2.0). (**GSE136685_AdditionalFile6_DatasetS5.txt**)

**Additional file 7: Dataset S6**. Normalized expression levels (TPM) of PCGs, binned by expression group. (**Additional_file7_DatasetS6.xlsx**)

**Additional file 8: Dataset S7**. Functional annotations, differential accessibility and expression data for differentially regulated PCGs in tassel vs. ear. (**Additional_file8_DatasetS7.xlsx**)

**Additional file 9: Dataset S8**. Genes bound by FEA4 and/or KN1 TFs in MNase HS regions of their proximal promoters, and functional enrichment using GO. (**Additional_file9_DatasetS8.xlsx**)

**Additional file 10: Dataset S9**. Distance matrix between ABI3 motifs and FEA4 binding sites, closest gene models and annotations associated with these combinatorial *cis*-element modules, and functional enrichment analysis using maize-GAMER GO for genes within 1kb of FEA4-ABI3 cis-element modules. (**Additional_file10_DatasetS9.xlsx**)

**Additional file 11: Dataset S10**. Tassel and ear footprints at the iSeg threshold (iSeg bc 4.0). (**GSE136685_Addition alFile11_DatasetS10.txt**)

**Additional file 12: Dataset S11**. Annotated high-confidence lncRNAs from inflorescence primordia (genomic coordinates, sequences and expression levels). (**Additional_file12_DatasetS11.xlsx**)

**Additional file 13: Dataset S12**. High-confidence lncRNAs supported by Sanger sequencing. (**Additional_file13_DatasetS12.txt**)

**Additional file 14: Dataset S13**. Classification of inflorescence-expressed high-confidence lncRNAs. (**Additional_file14_DatasetS13.xlsx**)

**Additional file 15: Dataset S14**. SNP-trait associations and closest gene annotations from GWAS analyses of TBN and ERN using all HapMap V3 SNPs and a subset within MNase HS regions only. (**Additional_file15_DatasetS14.xlsx**)

**Additional file 16: Dataset S15**. Output from stepwise GWAS analysis of TBN and ERN using both HapMapv3 SNPs and markers subset based on MNase HS regions. (**GSE136685_AdditionalFile16_DatasetS15.zip.gz**)

**Additional file 17**. Review history.

**Additional file 1. Supplemental figures and tables:**

**Figure S1**. Overview of biological samples, nuclei isolation, and titration of MNase concentration used in this study.

**Figure S2**. Comparison of peak-calling algorithms and biological stringencies with respect to *in planta* TF binding data.

**Figure S3.** Locus-specific comparison of MNase-seq coverage maps across biological replicates.

**Figure S4**. Distribution of tassel- and ear-specific HS sites across genomic features.

**Figure S5**. MNase read density is differentially enriched in 5’ and 3’ regions of genes when comparing DNS (light – heavy) and small fragment profiles.

**Figure S6**. MNase HS associated with transposable elements in maize inflorescence primordia.

**Figure S7**. MNase hypersensitivity compared among genes with varying expression levels in ear primordia.

**Figure S8**. Classification of TFs by family that were differentially regulated between tassel and ear primordia.

**Figure S9**. Analysis of chromatin accessibility data from ear primordia in the context of KN1 and FEA4 occupancy (parallel analyses from tassel presented in Figure 3).

**Figure S10**. *De novo* motif discovery within MNase HS regions that overlap annotated FEA4 binding sites.

**Figure S11**. Test for association of FEA4 binding sites with MNase HS footprints.

**Figure S12**. Pipeline for annotating and quantifying expression of lncRNAs.

**Figure S13**. Genic features of lncRNAs.

**Figure S14**. Spatiotemporal expression of lncRNAs in developing inflorescence primordia.

**Figure S15**. Co-expression analysis between lncRNAs and protein-coding gene (PCG) pairs.

**Figure S16**. FDR-adjusted p-values for MNase only SNP sets compared to all markers using GWAS with a small diversity panel.

**Table S1**. Summary statistics for MNase-seq data.

**Table S2**. MNase sensitive iSeg calls (DNS) using all fragments at different biological cutoffs.

**Table S3**. MNase sensitive iSeg calls using LCS at different biological cutoffs.

**Table S4**. Distribution of the MNase HS regions (bc 2.0) across genomic features.

**Table S5**. *De novo* motif enrichment analysis in footprints proximal to differentially regulated genes between tassel and ear.

**Table S6**. Average density of the experimentally validated TCP PWMs in tassel and ear footprints.

