## Additional File 1 for "The regulatory landscape of early maize inflorescence development"

**Supplemental figures**

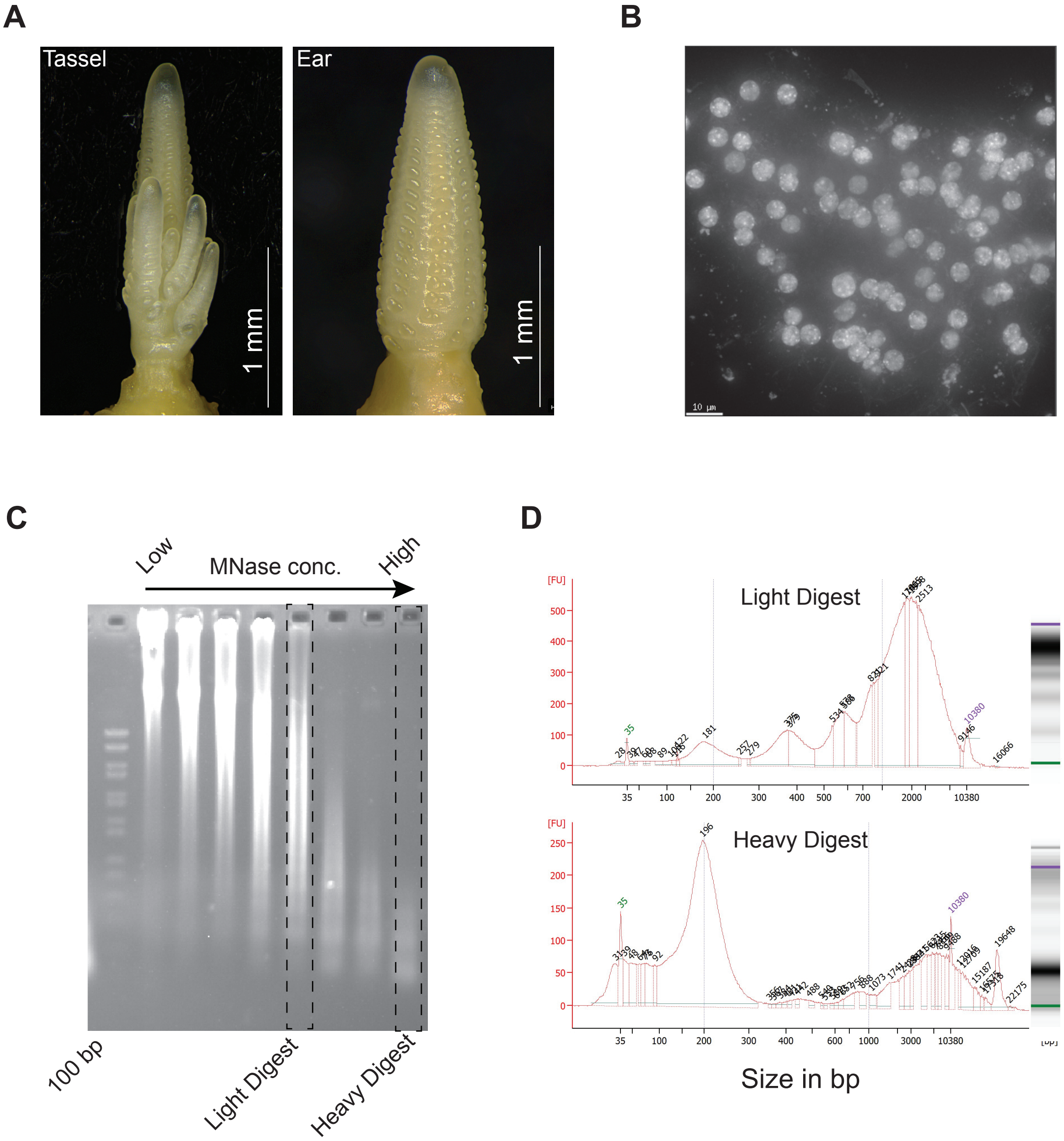

**Figure S1. Overview of biological samples, nuclei isolation, and titration of MNase concentration used in this study. A.** Representative images of hand-dissected tassel and ear primordia. **B.** Representative image of fixed, purified nuclei stained with DAPI from inflorescence primordia. **C.** Analytical gel of the DNA following digestion with varying concentrations of MNase. Digest levels selected for "light" and "heavy" MNase libraries are indicated. **D.** Agilent Bioanalyzer electropherogram traces of the light and heavy MNase digested DNA are shown alongside corresponding densitometry plots.

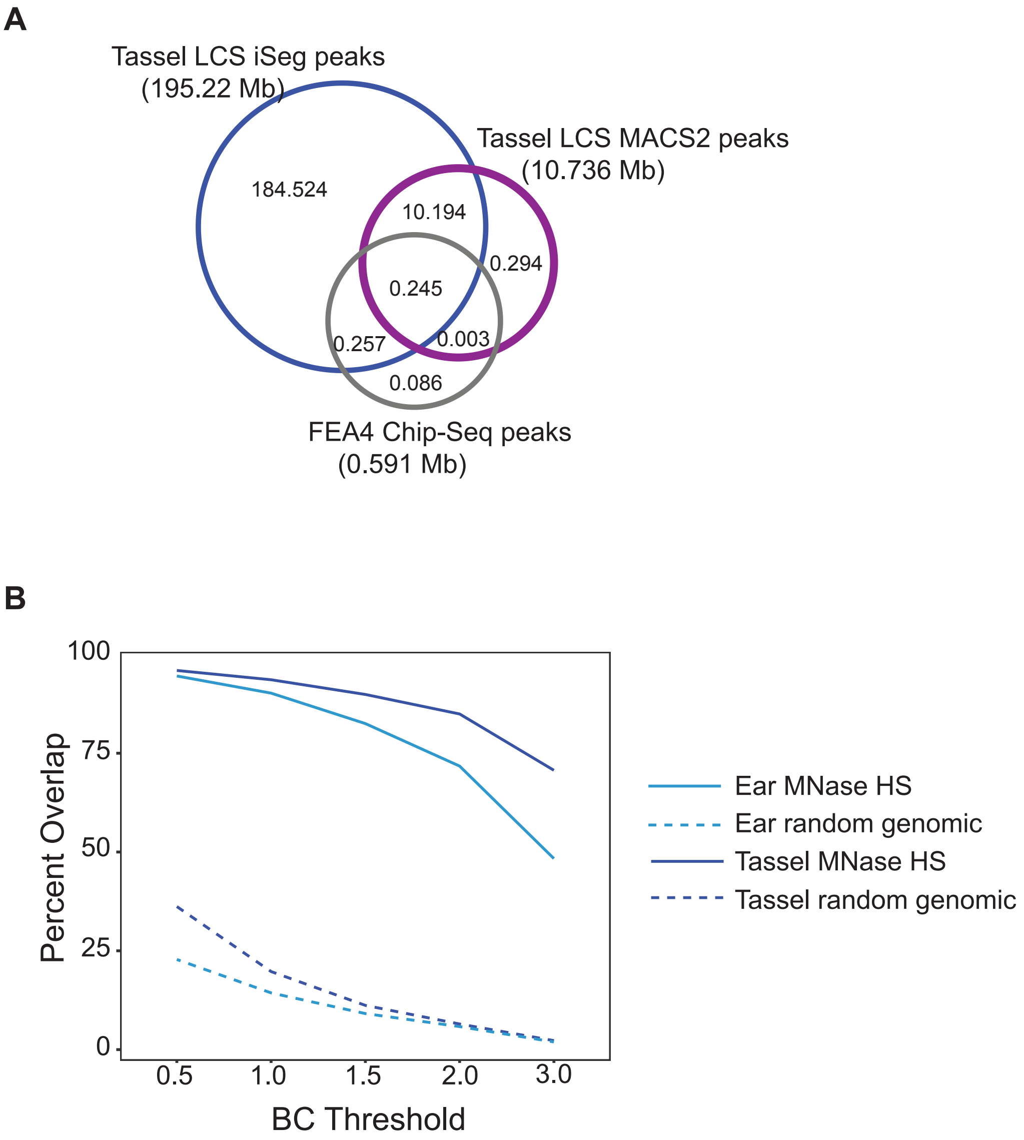

**Figure S2. Comparison of peak-calling algorithms and biological stringencies with respect to *in planta* TF binding data. A.** MNase HS peaks called by iSeg (bc 2.0) overlapped a larger fraction of FEA4 binding sites from ChIP-seq analyses in tassel primordia compared with MACS2. MNase HS regions are based on analysis of small sized fragments (<131bp) from the light digest (LCS). **B.** Total bp overlap of high confidence FEA4 ChIP-seq peaks with MNase HS regions called from different peak-calling stringencies in iSeg showed that decreasing stringency from bc 3.0 to 2.0 increased the overlap with TF-bound regions but did not significantly increase noise. Overlaps with TF binding were compared with a random set of genomic regions of equal size (shuffled at 100 repetitions) at each biological cutoff.

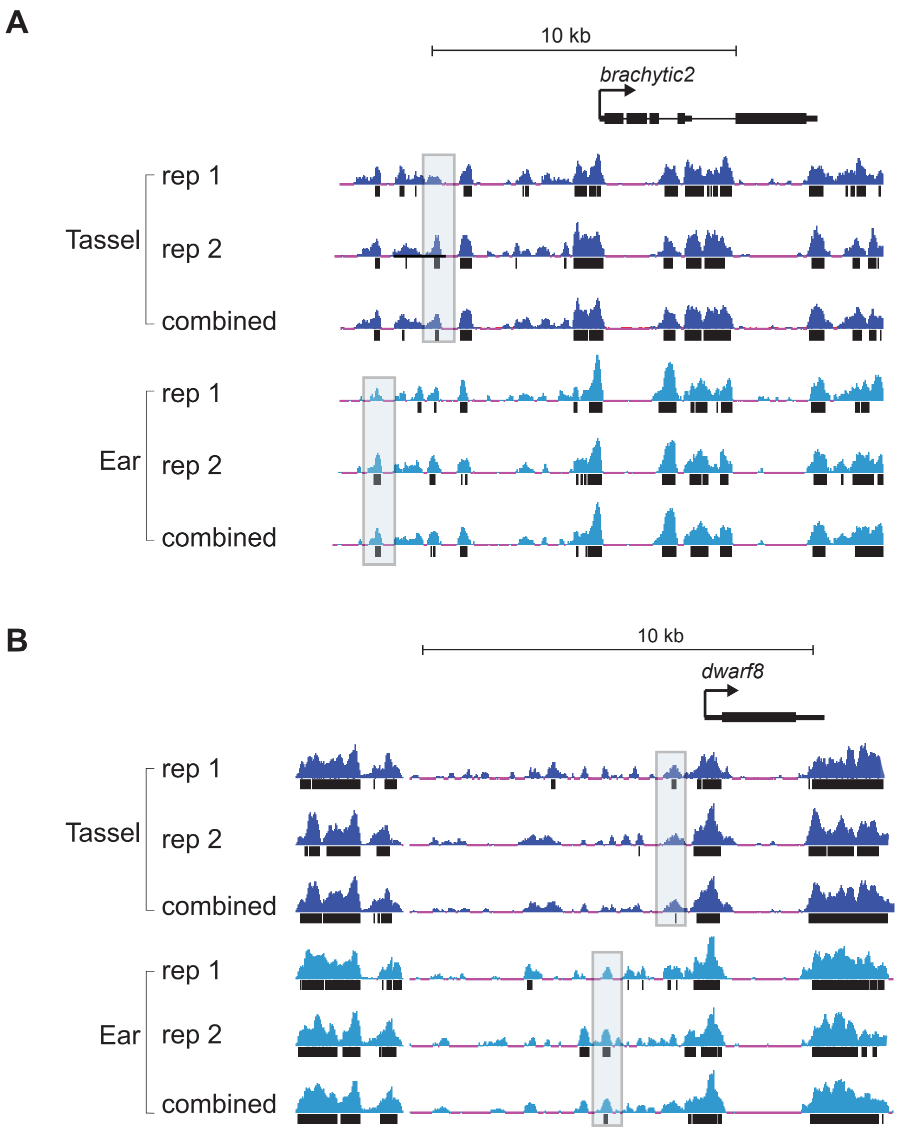

**Figure S3. Locus-specific comparison of MNase-seq coverage maps across biological replicates.** Genome browser views of MNase HS (DNS analysis) comparing individual biological replicates and a combined replicate track surrounding the **A**. *brachytic2* and **B**. *dwarf8* genes in tassel and ear primordia. Highlighted boxes indicate regions called as HS in combined replicate track, but not in one of the biological replicates due to coverage.

A

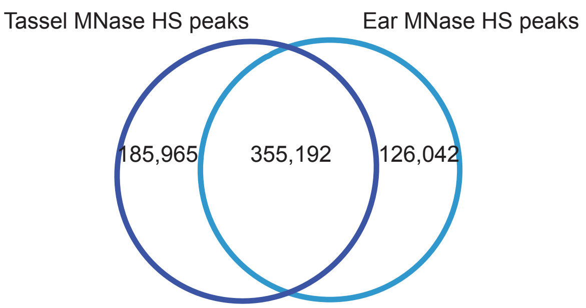

B

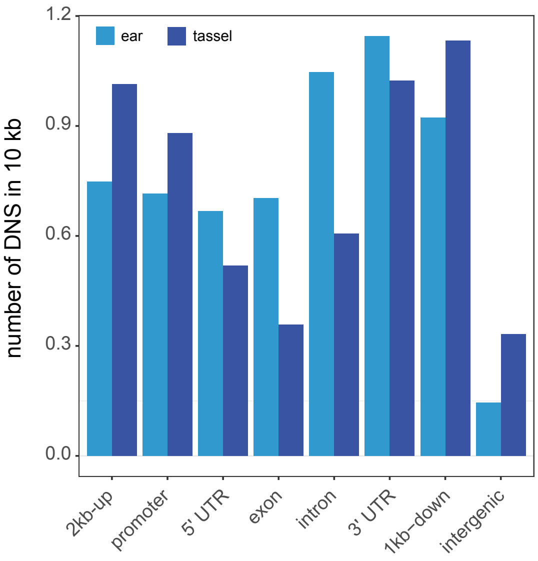

**Figure S4. Distribution of tassel- and ear-specific HS sites across genomic features. A**.Venn diagram compares unique and shared DNS sites mapped between tassel and ear primordia. **B.** Distribution of unique DNS sites from tassel and ear across genomic features shows organ-specific accessibility preferences

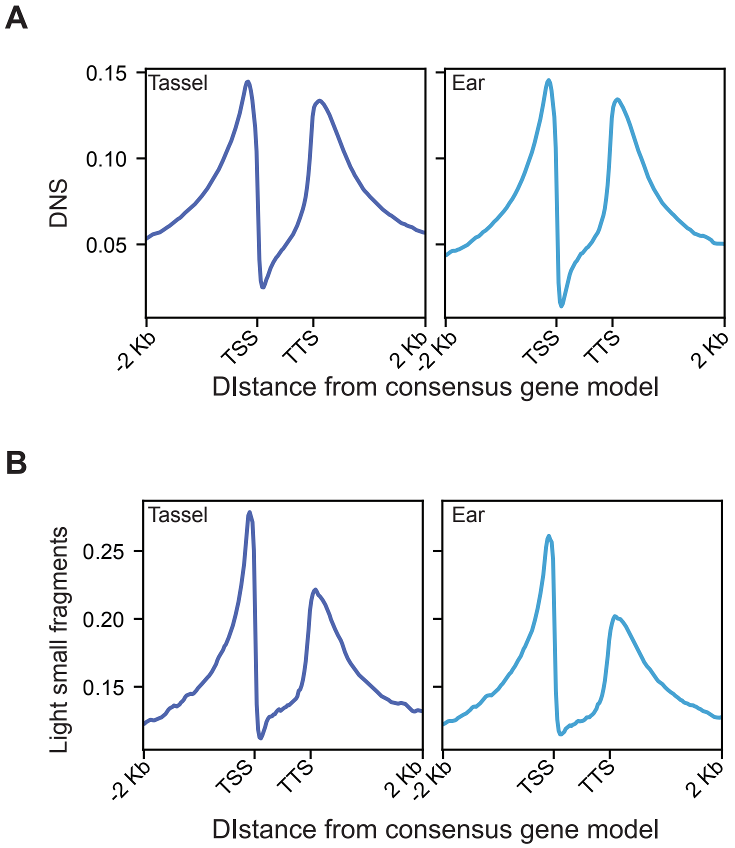

**Figure S5. MNase read density is differentially enriched in 5' and 3' regions of genes when comparing DNS (light - heavy) and small fragment profiles.** Distributions of MNase read density across a consensus gene model based on 39,151 protein-coding genes and their 2 kb flanking sequences are shown for **A.** DNS (light - heavy) and **B.** small fragments (< 131 bp) from the light MNase digest. Both profiles show enrichment of MNase HS in the promoter region. An almost equal enrichment is observed at the 3' end in the DNS profiles for both tassel and ear, which is markedly reduced in the small fragment profiles. This is likely indicative of the DNS analyses providing information on nucleosome occupancy in addition to TF binding, where the small fragment data aligns with binding of transcriptional complexes.

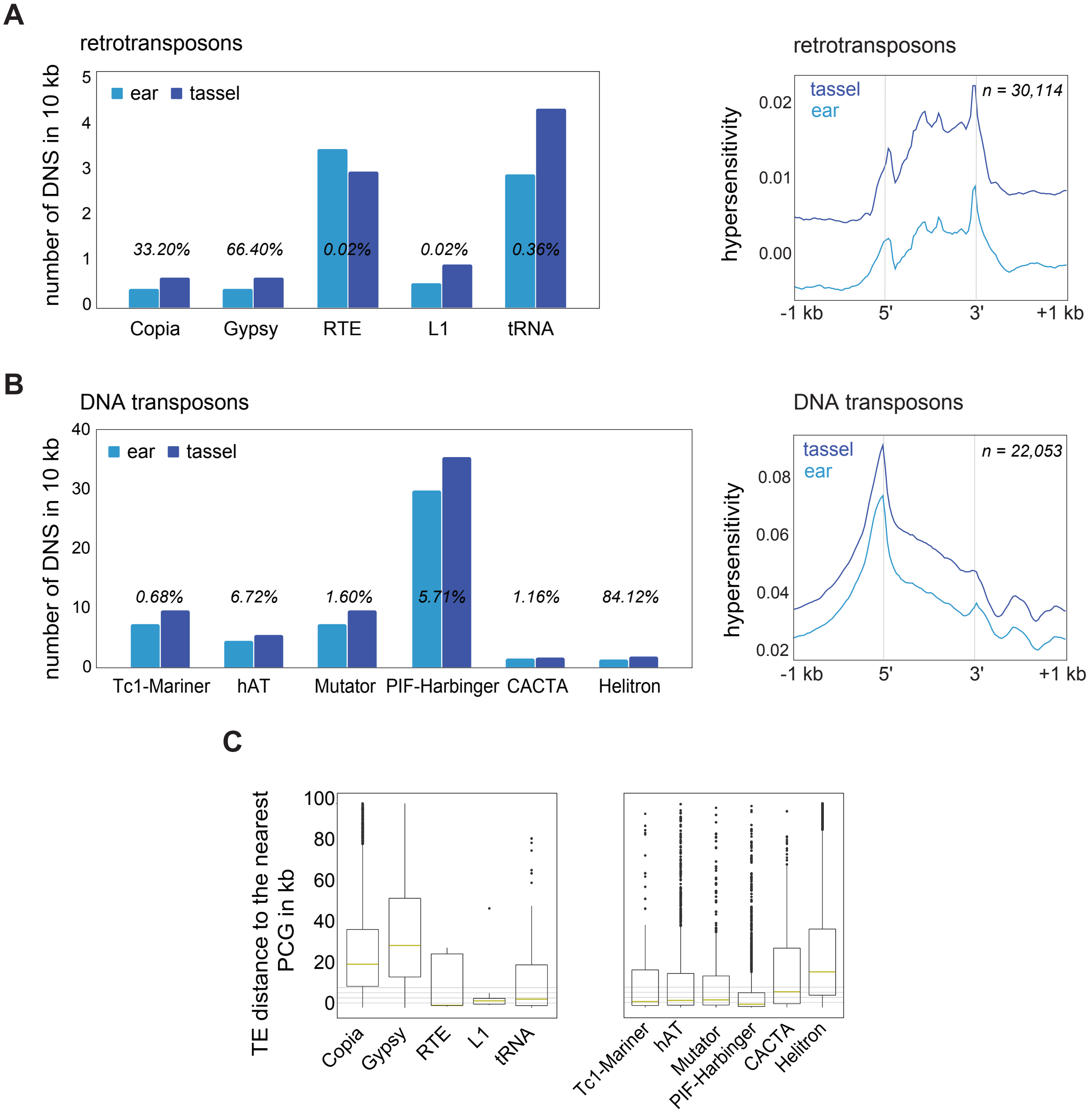

**Figure S6. MNase HS associated with transposable elements in maize inflorescence primordia.** **A.** Left panel: Number of intergenic MNase HS midpoint sites (bc 2.0) overlapping TE coordinates divided by retrotransposon class. Frequencies of MNase HS sites were normalized per 10 kb of genomic space per TE superfamily. Percent of each class represented in right panel are shown. Right panel: MNase HS plotted as average read density DNS (bc 2.0) at ± 1,000 bp surrounding the 5' and 3' ends of annotated retrotransposons. LTR (Copia, Gypsy); LINE (RTE, L1); SINE (tRNA). **B.** Left panel: Number of intergenic MNase HS midpoint sites (bc 2.0) overlapping TE coordinates divided by DNA transposon class. Frequencies of MNase HS sites were normalized per 10 kb of genomic space per TE superfamily. Right panel: MNase HS plotted as average read density DNS (bc 2.0) at ± 1,000 bp surrounding the 5' and 3' ends of annotated DNA transposons. TIR (Tc1-Mariner, hAT, Mutator, PIF-Harbinger, CACTA). **C.** Distance of accessible TEs to the closest PCGs; the yellow line represents the median distance.

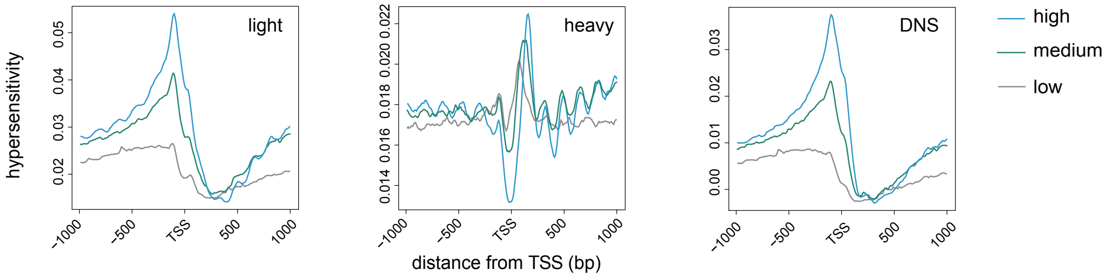

**Figure S7. MNase hypersensitivity compared among genes with varying expression levels in ear primordia.** 39,151 protein-coding genes were divided into 3 tertiles (high, medium and low expression) based on their TPM values. For each of the three groups, average accessibility and nucleosome occupancy was plotted at ± 1,000 bp surrounding the TSS. Lines represent the average read density of light and heavy MNase digests as well as the differential (light -heavy; DNS).

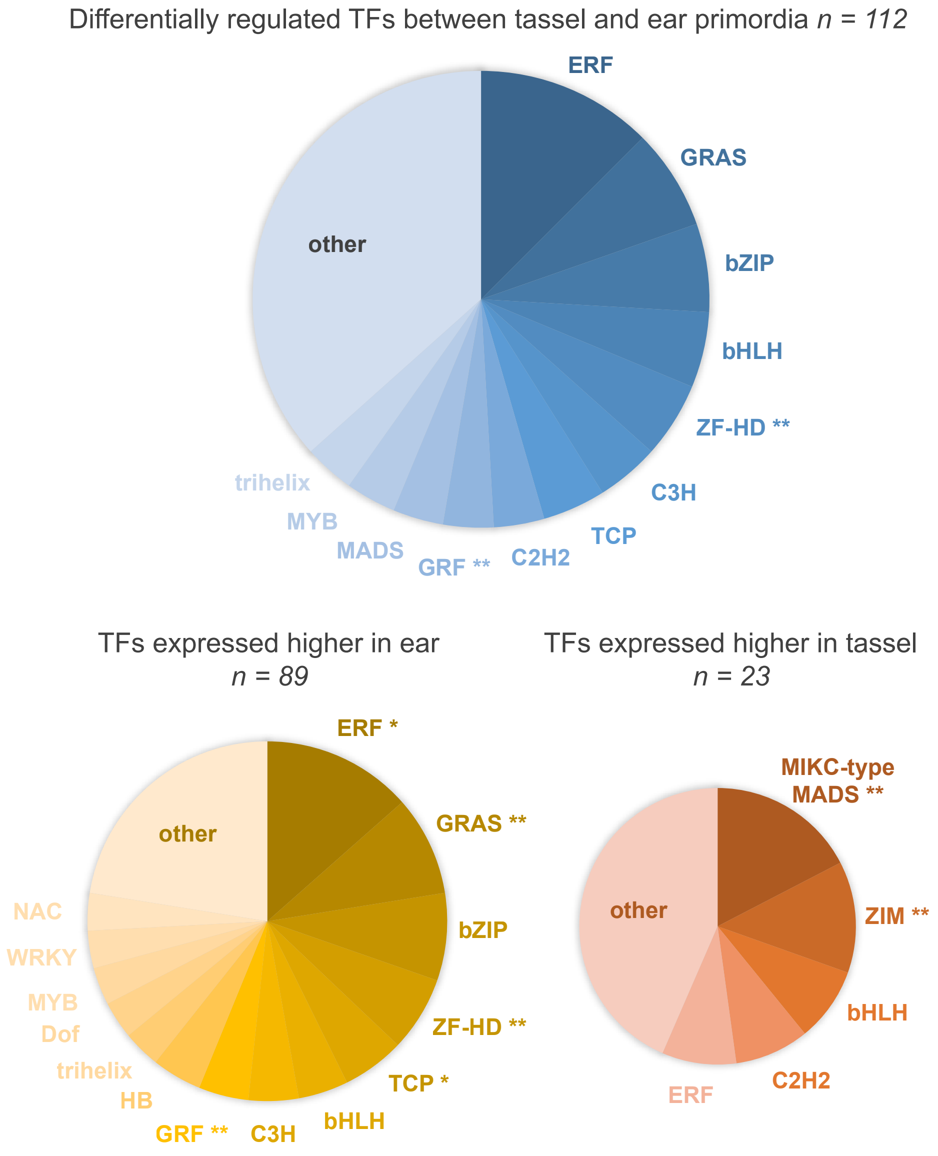

**Figure S8. Classification of TFs by family that were differentially regulated between tassel and ear primordia.** Classification of TFs by family that were differentially regulated between tassel and ear primordia. There is a clear shift in the expression of certain TF families between tassel and ear, suggesting these families are likely involved in meristem determinacy programs or early signaling of male- and female-specific floral development. TF families that were significantly enriched based on hypergeometric test within respective sets are indicated (** $q < 0.01$; * $q \leq 0.05$).

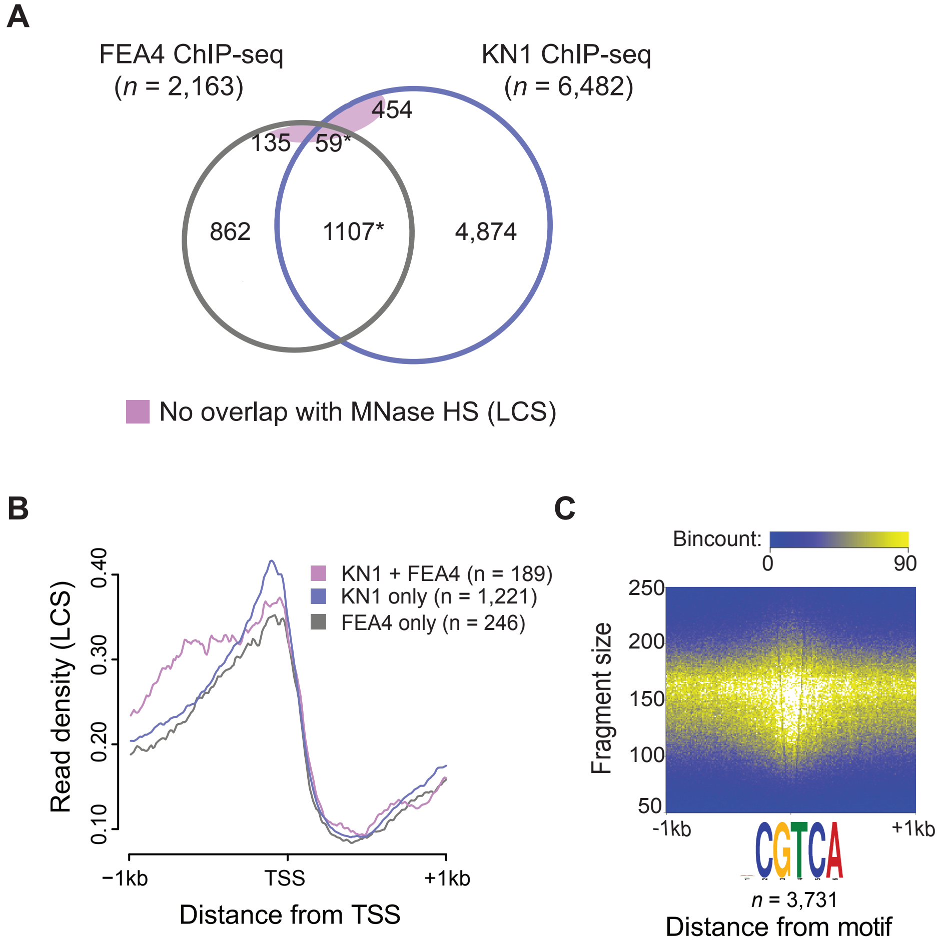

**Figure S9. Analysis of chromatin accessibility data from ear primordia in the context of KN1 and FEA4 occupancy (parallel analyses in tassel presented in Figure 3). A.** Total number of high confidence FEA4 and KN1 ChIP-seq peaks that overlap MNase HS regions from ear. * represents the number of FEA4 peaks shared with KN1. **B.** Read density (reads per million) of LCS centered around the TSSs of genes bound by KN1, FEA4 or co-bound by both TFs in their proximal promoters. **C.** Density of fragments from MNase light digests at FEA4 binding sites with consensus motif.

**A**

| Motifs | E-value | Closest matches (p-value) |
|---|---|---|
| 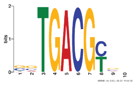 | 1.9e-250 | TGA1 (8.18e-06), TGA7 (1.11e-05)[a] |
| 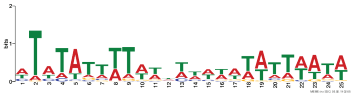 | 4.6e-013 | ATHB23 (6.82e-06), OBP3 (1.12e-05) |
| 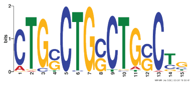 | 3.1e-008 | ERF6 (4.64e-03), ABI3 (6.02e-03)[b] |
| 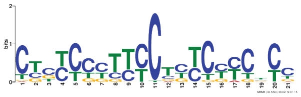 | 1.2e-005 | BPC1 (1.08e-06), BPC6 (1.88e-06) |

**B**

| Motifs | E-value | Closest matches (p-value) |
|---|---|---|
| 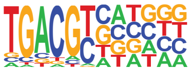 | 1e-1049 | bZIP910 (1.73e-04), TGA2(2.71e-04)[a] |
| 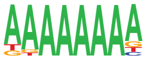 | 1e-191 | OBP3 (5.10e-05), AT5G66940 (7.12e-05) |
| 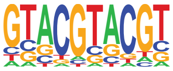 | 1e-131 | SPL11 (4.33e-03), SPL4 (4.59e-03) |
| 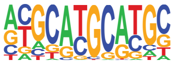 | 1e-128 | ABI3 (7.53e-03), FUS3 (2.66e-03) |
| 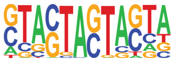 | 1e-116 | STZ (8.08e-03) |
| 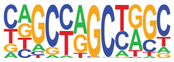 | 1e-109 | bZIP52 (6.30e-03), MYB3 (1.43e-02) |
| 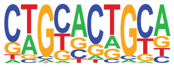 | 1e-103 | AT5G18450 (8.17e-03), AT5G18450 (8.47e-03) |
| 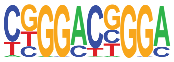 | 1e-99 | POPTR_0002s00440g (1.69e-03), SPL1 (4.75e-03)[c] |
| 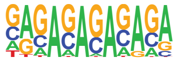 | 1e-98 | BPC6 (4.67e-10), ramosa1 (4.81e-10) |
| 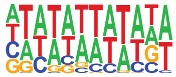 | 1e-94 | AGL3 (3.73e-03), AHL12 (3.77e-03) |

**Figure S10.** ***De novo*** **motif discovery within MNase HS regions that overlap annotated FEA4 binding sites. A.** Position Weight Matrices (PWMs) that were identified as significantly enriched using MEME, and closest matches to experimentally validated PWMs in the JASPAR Plant database. **B.** Top ten most significantly enriched PWMs (out of 26 total) identified using HOMER, and closest matches to experimentally validated PWMs in the JASPAR Plant database. Statistical significance of *de novo* motif enrichment is represented by E-values. The statistical significance of similarity between the de novo motif and the top two matches to experimentally validated PWMs in JASPAR are reported as p-values.

[a]The most significantly enriched motifs from both methods match either the FEA4 PWM exactly or similar bZIP PWM. [b]Manual inspection suggests that this de novo motif most closely matches the ABI3 PWM. [c]This motif closely matches the KN1 PWM in the HOMER database (KN1 and FEA4 PWMs were not present within the JASPAR Plant database).

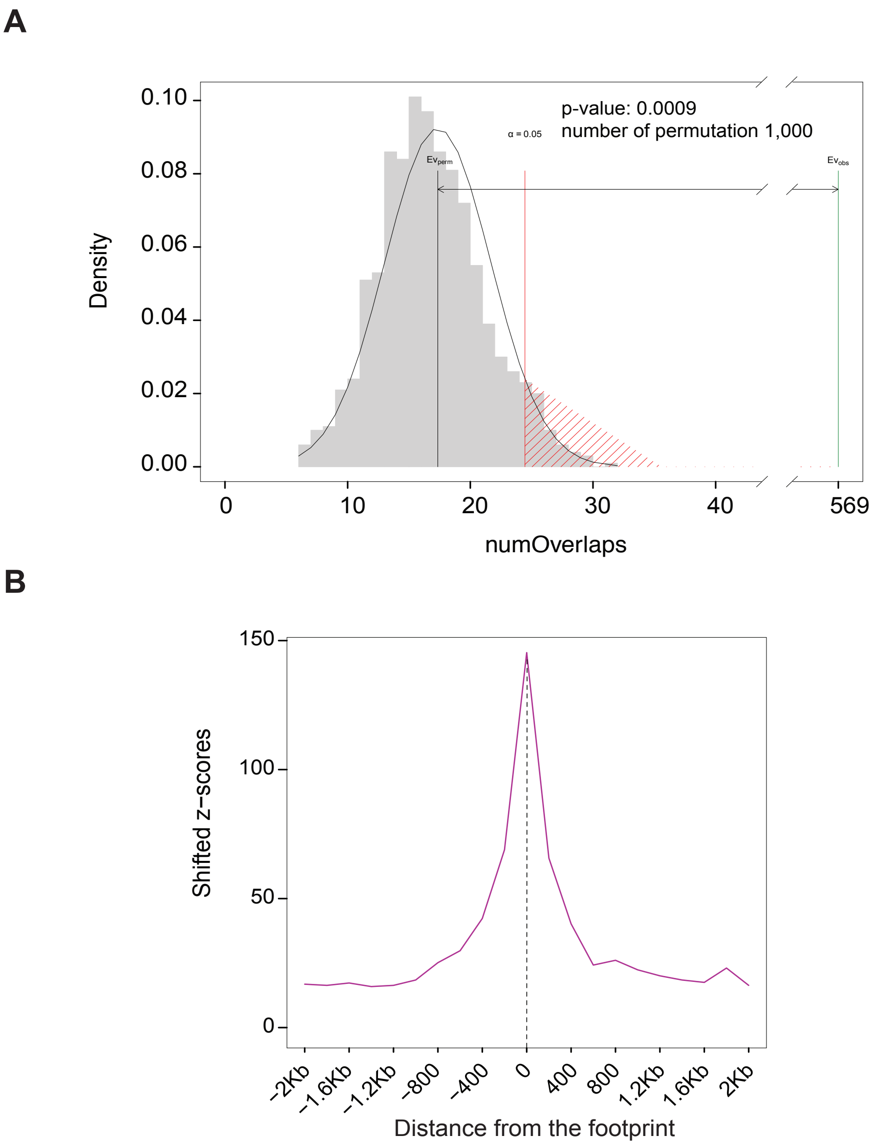

**Figure S11. Test for association of FEA4 binding sites with MNase HS footprints. A.** Representation of the permutation test (1,000 permutations) for the association of FEA4 binding sites with tassel MNase HS footprints. The grey area shows the distribution of number of overlaps among randomized regions. The black line represents the mean number of overlaps among randomized genomic regions (expected value), and the green line represents the number of overlaps between FEA4 binding sites and MNase HS footprints (observed value). The red line defines the significance limit. **B.** Distribution of Z-scores based on the permutation test in A indicates that the position of FEA4 binding sites is strongly dependent on the position of the MNase HS footprint. The Z-score markedly drops with small shifts of 100 bp from observed genomic positions.

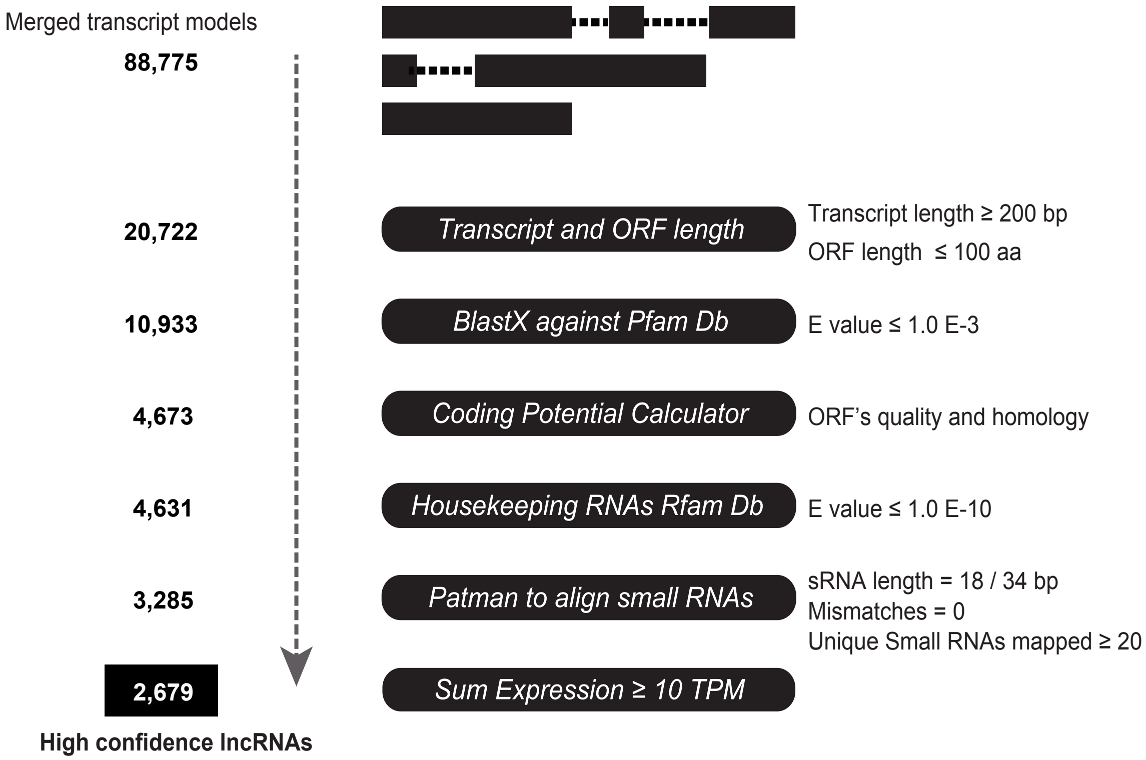

**Figure S12. Pipeline for annotating and quantifying expression of lncRNAs in maize inflorescence primordia.** Six consecutive filters were applied to identify high-confidence lncRNAs from a pool of reconstructed transcripts using a genome-guided approach starting from Illumina poly(A)+ RNA-seq data. These filters have been implemented taking into consideration the biochemical features of lncRNAs and current knowledge on lncRNA biology. This pipeline discriminates lncRNAs from protein-coding transcripts, structural RNAs, pre-miRNAs and all potential transcripts sharing perfect homology with small RNA sequences from young tassel and ear samples. To retain stringency without missing potential lncRNAs, two filters were applied: i) ORF size cut-off to retain transcripts encoding small peptides 100 amino acids (as described by Andrews et al. 2014, *Nature Reviews Genetics* 15 (3):193); ii) a sum expression cut-off of $\geq 10$ Transcript Per kilobase Million (TPM) across samples to remove transcripts of low expression. Numbers on the left represent transcripts retained after each filter, and cut-offs applied are specified on the right. 2,679 are the lncRNAs located on chromosomes 1 to 10 without including those annotated on contigs (*n*= 81).

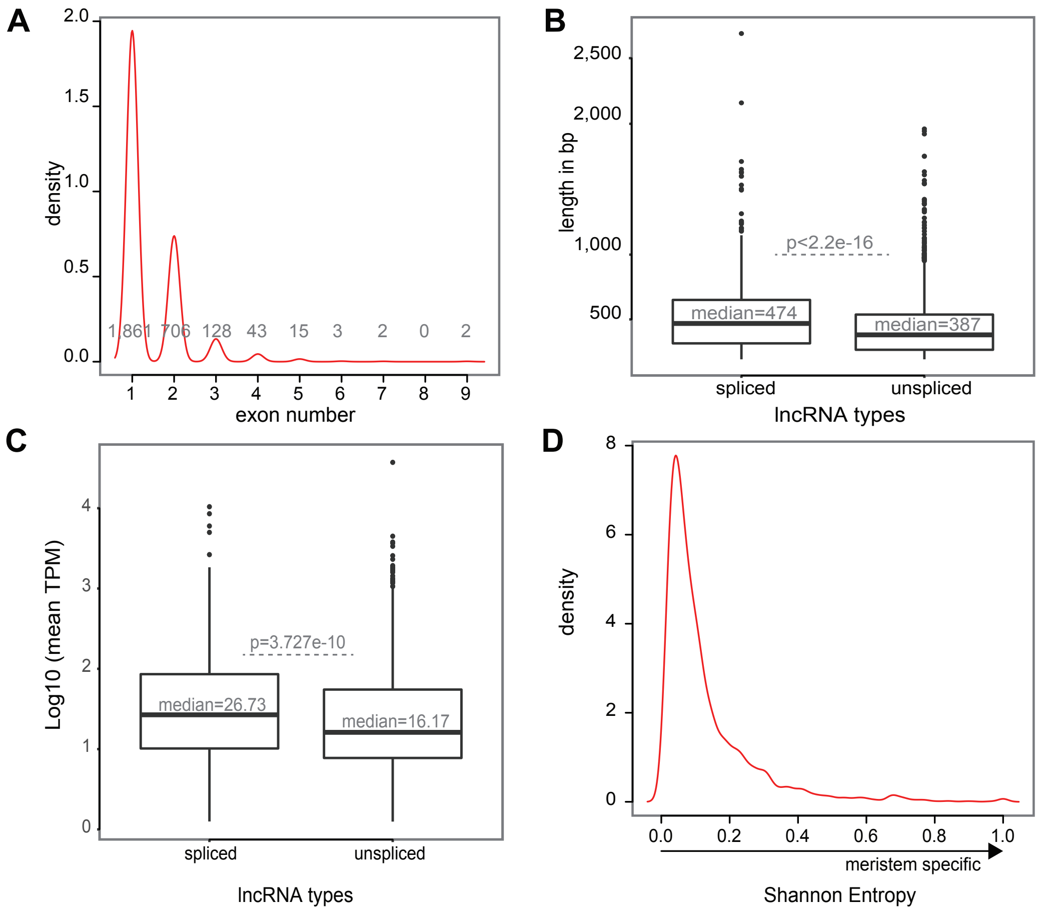

**Figure S13. Genic features of lncRNAs. A.** Exon number distribution among annotated lncRNAs. The x-axis represents exon number and y-axis represents the relative density of lncRNAs. The number below each peak represents the absolute number of lncRNAs belonging to each class. LncRNAs with 1 exon are defined as “unspliced”. **B.** Box-plot of transcript length for spliced and unspliced lncRNAs. **C.** Mean expression shown by a box-plot of maximum expression levels of spliced and unspliced lncRNAs. P-values in B and C were calculated using two-sided Wilcoxon test. **D.** To quantify spatiotemporal specific expression we used the Shannon Entropy (SH). SH was calculated starting from the TPM using the function entropy Specificity (option, norm=TRUE) written in the Bioconductor package BioQC. LncRNAs with spatiotemporal specific expression during early tassel and ear development were called by selecting SH $\geq$ 0.6 and assigned to the sample with the highest expression.

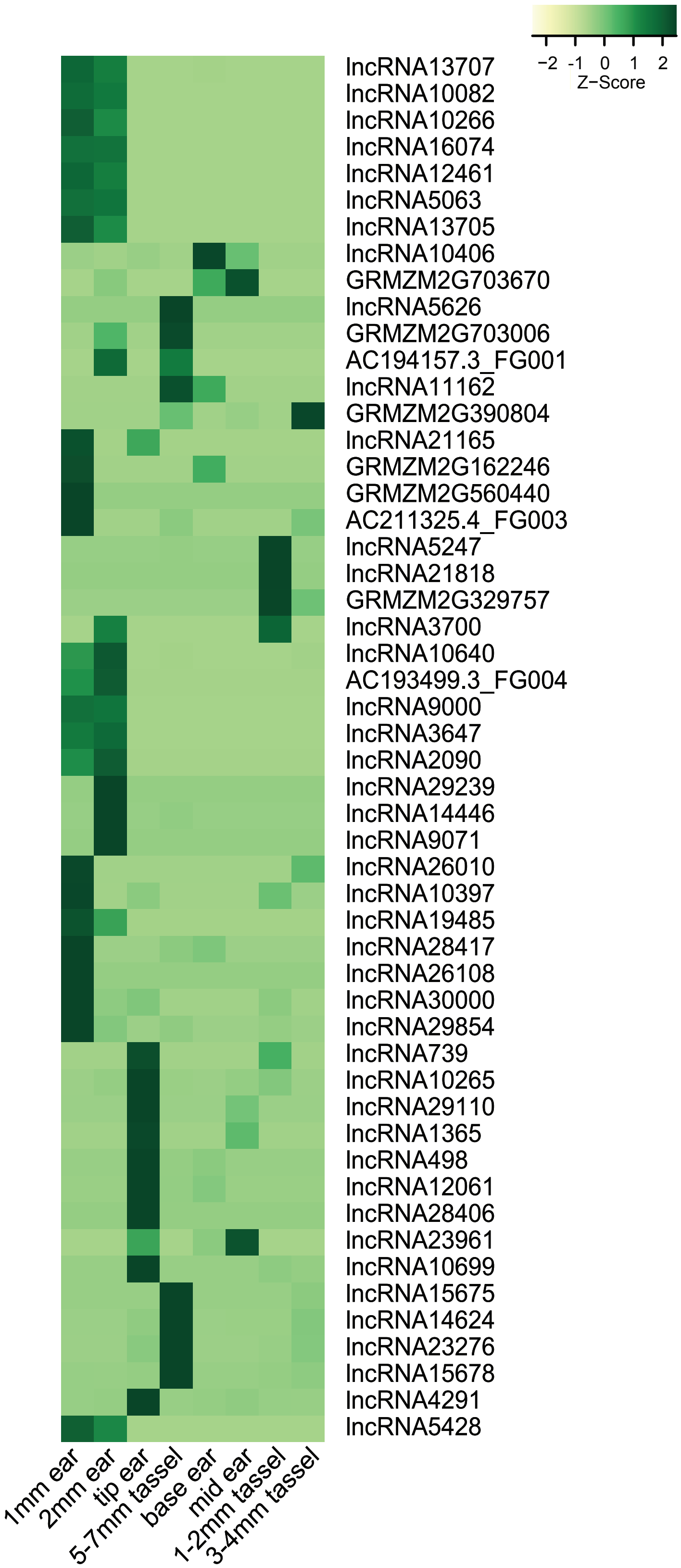

**Figure S14. Spatiotemporal expression of lncRNAs in developing inflorescence primordia.** The heatmap represents differential expression profiles of lncRNAs identified as showing spatiotemporal specificity in developing tassel and ear primordia based on Shannon Entropy (SH ≥ 0.6). Rows (lncRNAs) and columns (developmental stages and/or specific meristem types of tassel and ear primordia) were hierarchically clustered (Ward D method). Row-normalized expression level is presented as Z-score [(observed transcripts per million (TPM) – row mean TPM) / standard deviation of TPMs for that row]. Light yellow-to-dark green represents low-to-high expression with respect to the row mean.

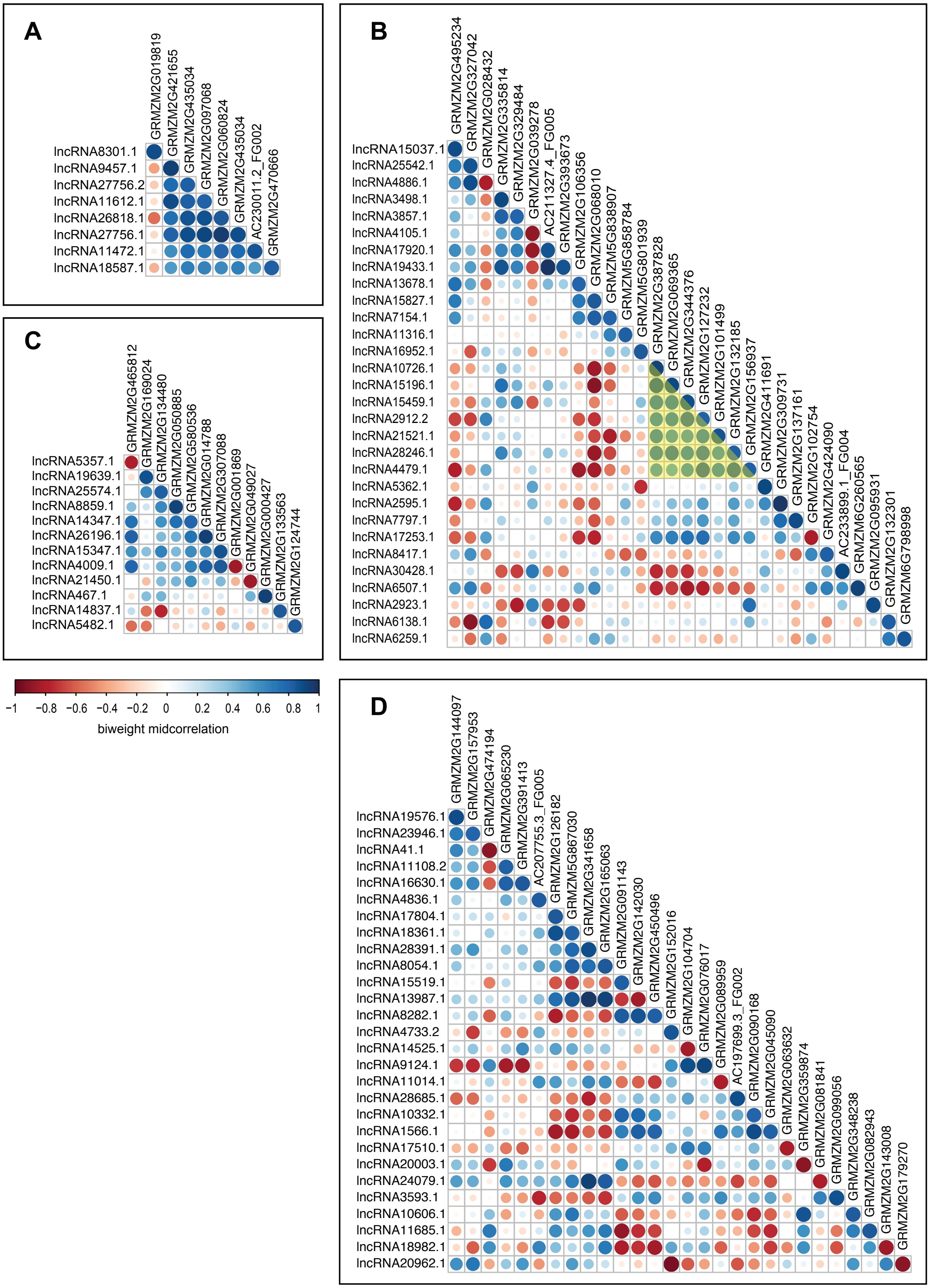

**Figure S15. Co-expression analysis between lncRNAs and protein-coding gene (PCG) pairs.** Co-expression profiles of lncRNAs and their closest PCGs across early tassel and ear development for: **A.** intronic-lncRNAs, **B.** intergenic-lncRNAs < 2 kb, **C.** intergenic-lncRNAs <10 kb. **D.** intergenic-lncRNAs >10 kb, Co-expressed pairs were determined using the biweight midcorrelation method and selecting for a coefficient > |0.8|. The selected pairs were then plotted and clustered based on the hierarchical clustering method using the R package *corrplot* allowing to highlight potential co-expressed modules among the lncRNA:PCG pairs. Correlated lncRNA-PCG pairs are shown in the diagonal of each image.

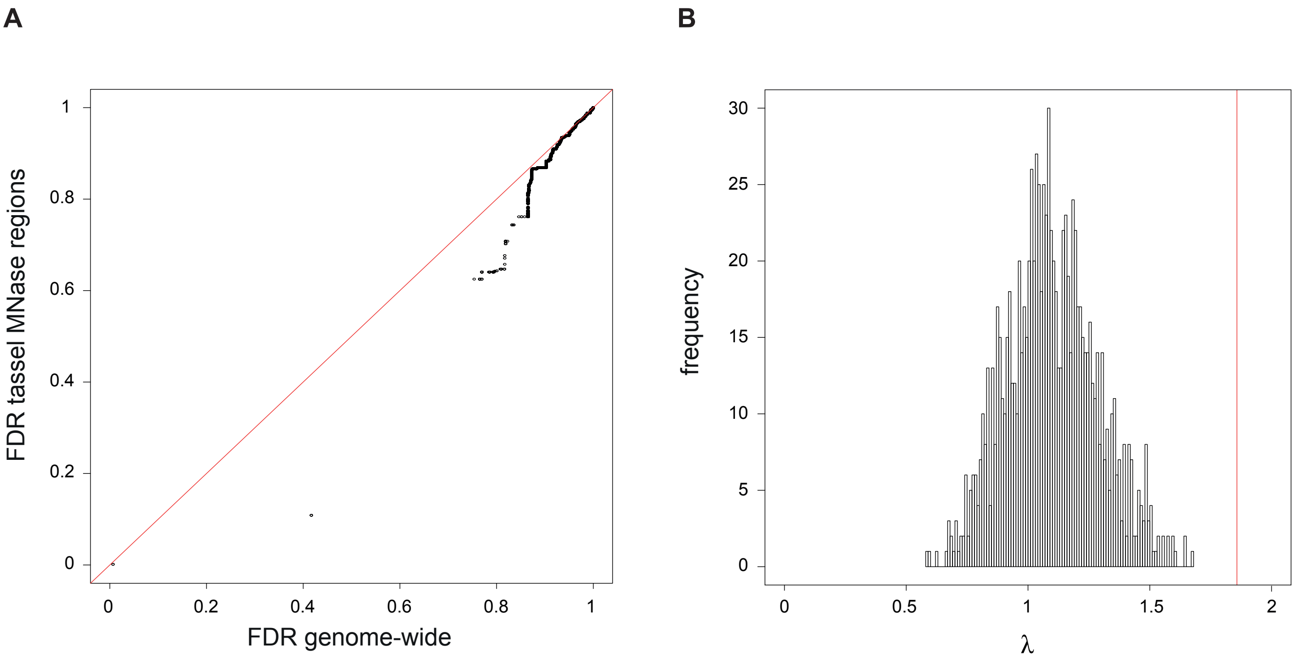

$$\lambda = \frac{\text{99th\_Percentile( -log(Reduced FDR adjusted p-values))}}{\text{99th\_Percentile(-log(Genome-wide FDR adjusted p-values))}}$$

**Figure S16. FDR-adjusted p-values for MNase-only SNP sets compared to all markers used in GWAS with a small diversity panel.** False Discovery Rate (FDR)-adjusted p-values (Benjamini and Hochberg, 1995) for TBN were used to compare significance of different marker subsets in GWAS.

**A)** Each point represents a SNP from a GWAS using the 282 lines maize diversity panel and public phenotype data for TBN (Rice et al., 2020). GWAS was run with 71,024 markers within tassel MNase regions and compared to the genome-wide marker set. Markers below the red line are more significant (have lower FDR-adjusted p-values) when only MNase regions were used compared to when the genome-wide marker set was used. The most significant loci from Rice et al. (2020) for TBN were present in tassel MNase regions. One marker on chromosome 10 showed a reduction from an adjusted p-value of 0.41 to 0.10 when only considering markers in tassel MNase regions.

**B)** The histogram distribution of lambda (λ) for 1,000 iterations of 71,024 randomly chosen markers, number of breaks = 100. λ is a measure of the decrease in the FDR adjusted p-values for the 99th percentile markers in the reduced marker set compared to the genome-wide marker set (equation). The vertical red line represents λ for markers in the tassel MNase regions. For the 71,024 markers in tassel MNase regions, the 99th percentile means that we picked the top 710 markers with the highest -log(FDR-adjusted p-values), which given the conservative nature of GWAS, is a large number of SNPs to investigate.

**Table S1.** Summary statistics for MNase-seq data.
[1] A minimum alignment quality score of 20 was applied.
[2] Number of mapped reads <131 bp.

| Tissue | MNase digest | Biological replicate | # sequenced reads | Mapped pairs after QC[1] | # small fragments[2] |
|---|---|---|---|---|---|
| Tassel | light | 1 | 158,486,776 | 83,883,777 | 11,960,681 |
| | | 2 | 131,075,290 | 70,030,688 | 9,413,425 |
| | heavy | 1 | 183,700,621 | 88,946,020 | 41,181,987 |
| | | 2 | 164,897,173 | 75,337,700 | 20,342,150 |
| Ear | light | 1 | 197,317,939 | 96,131,890 | 23,106,990 |
| | | 2 | 171,343,121 | 84,119,398 | 13,880,748 |
| | heavy | 1 | 128,835,468 | 66,139,739 | 13,799,902 |
| | | 2 | 158,523,648 | 81,014,951 | 13,224,726 |

**Table S2.** MNase sensitive iSeg calls (DNS) using all fragments at different biological cutoffs.
[1] MNase sensitive regions based on iSeg were identified using differential nucleosome sensitivity (DNS) between the light and heavy digests (light minus heavy).
[2] Segments are considered overlapped if their midpoints are within 300 bp.
[3] The total number and percentage of peaks in biological replicate 2 that overlap with replicate 1.
[4] Overlap between the iSeg with the MACS2 peak calls (replicates combined). Segments are considered overlapped if 50% of overlaps with MACS2 peak. Total number of tassel MACS2 peaks = 213,283 and total number of ear MACS2 peaks = 198,582.

| | **Total size of the MNase sensitive regions (Mb)** | | | **Total number of MNase sensitive regions[1]** | | | | |
|---|---|---|---|---|---|---|---|---|
| **Biological cutoff (bc)** | **Rep1** | **Rep2** | **Reps combined** | **Rep 1** | **Rep 2** | **Reps combined** | **Replicate overlap[2,3] (%)** | **MACS2 overlap[4] (%)** |
| **Tassel** | | | | | | | | |
| 0.5 | 460.28 | 464.05 | 471.15 | 1,894,624 | 1,639,763 | 1,415,211 | 1,248,402 (76.13) | 212,287 (99.5) |
| 1.0 | 297.93 | 298.59 | 296.6 | 1,350,875 | 1,194,935 | 999,106 | 906,417 (75.85) | 212,143 (99.57) |
| 1.5 | 192.13 | 192.1 | 189.47 | 930,443 | 834,890 | 716,373 | 642,560 (76.96) | 210,153 (98.93) |
| 2.0 | 121.52 | 121.83 | 119.74 | 688,989 | 638,542 | 560,823 | 496,432 (77.74) | 197,491 (93.20) |
| 3.0 | 40.52 | 41.11 | 40.27 | 380,288 | 372,652 | 347,199 | 267,062 (71.67) | 125,926 (59.12) |
| **Ear** | | | | | | | | |
| 0.5 | 455.54 | 459.18 | 464.52 | 2,033,613 | 1,902,342 | 1,566,511 | 1,347,074 (70.81) | 197,512 (99.51) |
| 1.0 | 296.52 | 296.66 | 293.69 | 1,360,875 | 1,264,010 | 995,431 | 893,861 (70.72) | 197,403 (99.41) |
| 1.5 | 189.43 | 189.35 | 185.8 | 844,359 | 802,744 | 639,966 | 581,786 (72.47) | 195,993 (98.70) |
| 2.0 | 119.62 | 119.57 | 117.04 | 594,939 | 581,790 | 481,234 | 443,968 (76.31) | 183,048 (92.18) |
| 3.0 | 40.63 | 40.62 | 39.75 | 345,801 | 343,898 | 314,280 | 265,668 (77.25) | 105,628 (53.19) |

**Table S3.** MNase sensitive iSeg calls using LCS at different biological cutoffs.
[1] MNase sensitive regions based on iSeg were identified using the small fragments (<131bp) from the light digest.
[2] Segments are considered overlapped if they are placed within 300bp of their midpoints
[3] The total number and percentage of peaks in rep2 that overlap with rep1.
[4] Overlap between the iSeg with the MACS2 peak calls (reps combined). Segments are considered overlapped if 50% of the iSeg overlaps with MACS2 peak or vice versa. Total number of tassel MACS2 peaks = 48,415 and total number of ear MACS2 peaks = 57,219.

| | Total size of the MNase sensitive regions (Mb) | | | Total number of MNase sensitive regions[1] | | | | |
|---|---|---|---|---|---|---|---|---|
| **Biological cutoff (bc)** | **Rep1** | **Rep2** | **Reps combined** | **Rep 1** | **Rep 2** | **Reps combined** | **Replicate overlap[2,3] (%)** | **MACS2 overlap[4] (%)** |
| **Tassel** | | | | | | | | |
| 0.5 | 796.89 | 670.98 | 827.12 | 3,449,429 | 3,465,881 | 2,410,903 | 2,995,547 (86.43) | 47,990 (99.12) |
| 1.0 | 484.69 | 407.52 | 505.41 | 2,247,026 | 1,959,769 | 2,037,466 | 1,563,251 (79.77) | 47,931 (99) |
| 1.5 | 264.19 | 299.14 | 318.31 | 1,430,434 | 2,385,106 | 1,551,298 | 1,539,485 (64.55) | 47,932 (99) |
| 2.0 | 205.17 | 162.81 | 195.22 | 1,618,603 | 1,237,751 | 1,143,006 | 964,115 (77.89) | 47,985 (99.11) |
| 3.0 | 65.49 | 67.77 | 73.66 | 506,819 | 690,310 | 607,540 | 383,844 (55.6) | 48,040 (99.23) |
| **Ear** | | | | | | | | |
| 0.5 | 716.90 | 793.41 | 749.47 | 2,577,234 | 3,407,064 | 2,356,007 | 2,539,587 (74.54) | 56,554 (98.84) |
| 1.0 | 430.56 | 425.40 | 410.12 | 2,376,589 | 2,649,207 | 1,598,867 | 2,033,077 (76.74) | 56,576 (98.88) |
| 1.5 | 223.10 | 247.72 | 229.61 | 1,073,473 | 1,596,294 | 1,115,158 | 999,639 (62.62) | 56,619 (98.95) |
| 2.0 | 136.42 | 142.00 | 134.94 | 902,239 | 1,054,399 | 838,557 | 731,433 (69.37) | 56,647 (99) |
| 3.0 | 47.96 | 51.93 | 46.38 | 403,668 | 444,107 | 353,448 | 295,391 (66.51) | 55,865 (97.63) |

**Table S4**. Distribution of the MNase HS regions (bc 2.0) threshold within the genomic features.
[1] The midpoints of the MNase HS regions were overlapped with the features.
[2] The proximal promoter is defined as 1kb upstream and 200bp downstream of the annotated TSS of the primary transcript.
[3] The intergenic regions include all regions excluding the primary transcipts and their regulatory regions (2 kb upstream and 1kb downstream).

| Type | Genomic features | Sum of bp | Ear overlapped[1] | Ear overlapped (normalized to 10 kb) | Tassel overlapped | Tassel overlapped (normalized to 10 kb) |
|---|---|---|---|---|---|---|
| DNS | Exon | 41,063,856 | 11,496 | 2.79 | 10,024 | 2.44 |
| DNS | Intron | 92,809,401 | 48,520 | 5.22 | 49,135 | 5.29 |
| DNS | 5' UTR | 6,831,402 | 2,749 | 4.02 | 2,809 | 4.11 |
| DNS | 3' UTR | 10,900,303 | 8,725 | 8.00 | 9,036 | 8.28 |
| DNS | Promoter 2Kb-Up | 78,298,000 | 39,227 | 5.00 | 42,526 | 5.43 |
| DNS | Proximal promoter[2] | 46,978,800 | 23,800 | 5.06 | 25,564 | 5.44 |
| DNS | 1Kb-Down | 39,148,000 | 27,254 | 6.96 | 28,989 | 7.40 |
| DNS | Intergenic[3] | 1,806,641,379 | 234,938 | 1.30 | 306,364 | 1.69 |
| LCS | Exon | 41,063,856 | 21,458 | 5.22 | 24,073 | 5.86 |
| LCS | Intron | 92,809,401 | 74,563 | 8.03 | 74,657 | 8.04 |
| LCS | 5' UTR | 6,831,402 | 5,439 | 7.96 | 5,990 | 8.76 |
| LCS | 3' UTR | 10,900,303 | 13,049 | 11.97 | 12,345 | 11.32 |
| LCS | Promoter 2Kb-Up | 78,298,000 | 60,652 | 7.74 | 65,973 | 8.42 |
| LCS | Proximal promoter[2] | 46,978,800 | 36,217 | 7.70 | 37,623 | 8.00 |
| LCS | 1Kb-Down | 39,148,000 | 41,734 | 10.66 | 43,833 | 11.19 |
| LCS | Intergenic[3] | 1,806,641,379 | 450,449 | 2.49 | 737,501 | 4.08 |
| DNS specific | Exon | 41,063,856 | 2,889 | 0.70 | 1,470 | 0.35 |
| DNS specific | Intron | 92,809,401 | 9,717 | 1.04 | 5,628 | 0.60 |
| DNS specific | 5' UTR | 6,831,402 | 456 | 0.66 | 355 | 0.51 |
| DNS specific | 3' UTR | 10,900,303 | 1,248 | 1.14 | 1,116 | 1.02 |
| DNS specific | Promoter 2Kb-Up | 78,298,000 | 5,860 | 0.74 | 7,945 | 1.01 |
| DNS specific | Proximal promoter[2] | 46,978,800 | 3,359 | 0.71 | 4,135 | 0.88 |
| DNS specific | 1Kb-Down | 39,148,000 | 3,614 | 0.92 | 4,436 | 1.13 |
| DNS specific | Intergenic[3] | 1,806,641,379 | 26,284 | 0.14 | 60,062 | 0.33 |

**Table S5**. *De novo* motif enrichment analysis in footprints proximal to differentially regulated genes between tassel and ear.

[1] Footprints called using iSeg (bc4 threshold) were used in enrichment analyses for genes in two categories: showed differential changes in gene expression between tassel and ear and i) were more accessible in proximal promoters in tassel ($n$ = 383) or ii) ear ($n$ = 441).

[2] Closest match in the JASPAR Plant database using the motif comparison tool TOMTOM. The TF class that the closest match motif belongs to is provided in brackets.

| Category[1] | Motif PWM | Closest match[2] |
|---|---|---|
| > Ear accessibility | RGCCS | OsI_08196, TCP 15 [bHLH] |
| | CGSK | WRKY30 [WRKY] |
| | AGWGKG | AT3G46070 [C2H2 ZF] |
| | CACG | MYC4 [bHLH] |
| > Tassel accessibility | GGSS | ERF13 [AP2/ERF] |
| | CGSR | ARF3 [B3 domain] |

**Table S6**. Average density of experimentally validated TCP PWMs in footprints.
[1] Total occurrences of the PWM in the tassel and ear footprints scanned by FIMO.
[2] The average density normalized per kb in the tassel and ear footprints (Total bp of footprints in tassel = 9.97 Mbp and ear = 4.17 Mbp).

| Motif | Absolute count[1] | | Average density (per kb) [2] | | Ratio change[3] (Tassel/Ear) |
|---|---|---|---|---|---|
| | Ear | Tassel | Ear | Tassel | |
| PTF1 | 32,908 | 54,414 | 7.98 | 5.46 | 0.68 |
| TCP1 | 14,459 | 22,113 | 3.50 | 2.22 | 0.63 |
| TCP14 | 19,805 | 31,967 | 4.80 | 3.21 | 0.67 |
| TCP15 | 52,556 | 89,159 | 12.74 | 8.95 | 0.70 |
| TCP16 | 35,056 | 61,061 | 8.50 | 6.13 | 0.72 |
| TCP17 | 38,953 | 65,742 | 9.44 | 6.60 | 0.70 |
| TCP19 | 38,045 | 67,671 | 9.22 | 6.79 | 0.74 |
| TCP2 | 22,546 | 37,578 | 5.46 | 3.77 | 0.69 |
| TCP20 | 44,219 | 74,991 | 10.72 | 7.52 | 0.70 |
| TCP23 | 55,221 | 93,484 | 13.39 | 9.38 | 0.70 |
| TCP24 | 39,056 | 64,629 | 9.47 | 6.48 | 0.68 |
| TCP3 | 33,500 | 54,408 | 8.12 | 5.46 | 0.67 |
| TCP4 | 48,743 | 83,970 | 11.81 | 8.42 | 0.71 |
| TCP5 | 33,756 | 60,870 | 8.18 | 6.11 | 0.75 |
| TCP7 | 29,660 | 52,750 | 7.19 | 5.29 | 0.74 |
| At1g72010 | 33,050 | 55,618 | 8.01 | 5.58 | 0.70 |
| At2g45680 | 19,080 | 28,265 | 4.62 | 2.84 | 0.61 |
| At5g08330 | 31,630 | 51,339 | 7.67 | 5.15 | 0.67 |
| MYB27 | 18,861 | 37,584 | 4.57 | 3.77 | 0.82 |
| MYB57 | 19,320 | 37,903 | 4.68 | 3.80 | 0.81 |
| MYB62 | 19,673 | 37,239 | 4.77 | 3.74 | 0.78 |
